# Exploring G and C-quadruplex structures as potential targets against the severe acute respiratory syndrome coronavirus 2

**DOI:** 10.1101/2020.08.19.257493

**Authors:** Efres Belmonte-Reche, Israel Serrano-Chacón, Carlos Gonzalez, Juan Gallo, Manuel Bañobre-López

**Affiliations:** Advanced (magnetic) Theranostic Nanostructures Lab, INL-International Iberian Nanotechnology Laboratory, Av. Mestre José Veiga, 4715-330 Braga, Portugal; Instituto de Química Física ‘Rocasolano’, CSIC, 28006 Madrid, Spain

## Abstract

In this paper we report the analysis of the 2019-nCoV genome and related viruses using an upgraded version of the open-source algorithm G4-iM Grinder. This version improves the functionality of the software, including an easy way to determine the potential biological features affected by the candidates found. The quadruplex definitions of the algorithm were optimized for 2019-nCoV. Using a lax quadruplex definition ruleset, which accepts amongst other parameters two residue G- and C-tracks, hundreds of potential quadruplex candidates were discovered. These sequences were evaluated by their *in vitro* formation probability, their position in the viral RNA, their uniqueness and their conservation rates (calculated in over three thousand different COVID-19 clinical cases and sequenced at different times and locations during the ongoing pandemic). These results were compared sequentially to other *Coronaviridae* members, other Group IV (+)ssRNA viruses and the entire realm. Sequences found in common with other species were further analyzed and characterized. Sequences with high scores unique to the 2019-nCoV were studied to investigate the variations amongst similar species. Quadruplex formation of the best candidates was then confirmed experimentally. Using NMR and CD spectroscopy, we found several highly stable RNA quadruplexes that may be suitable theranostic targets against the 2019-nCoV.

**GRAPHICAL ABSTRACT:** 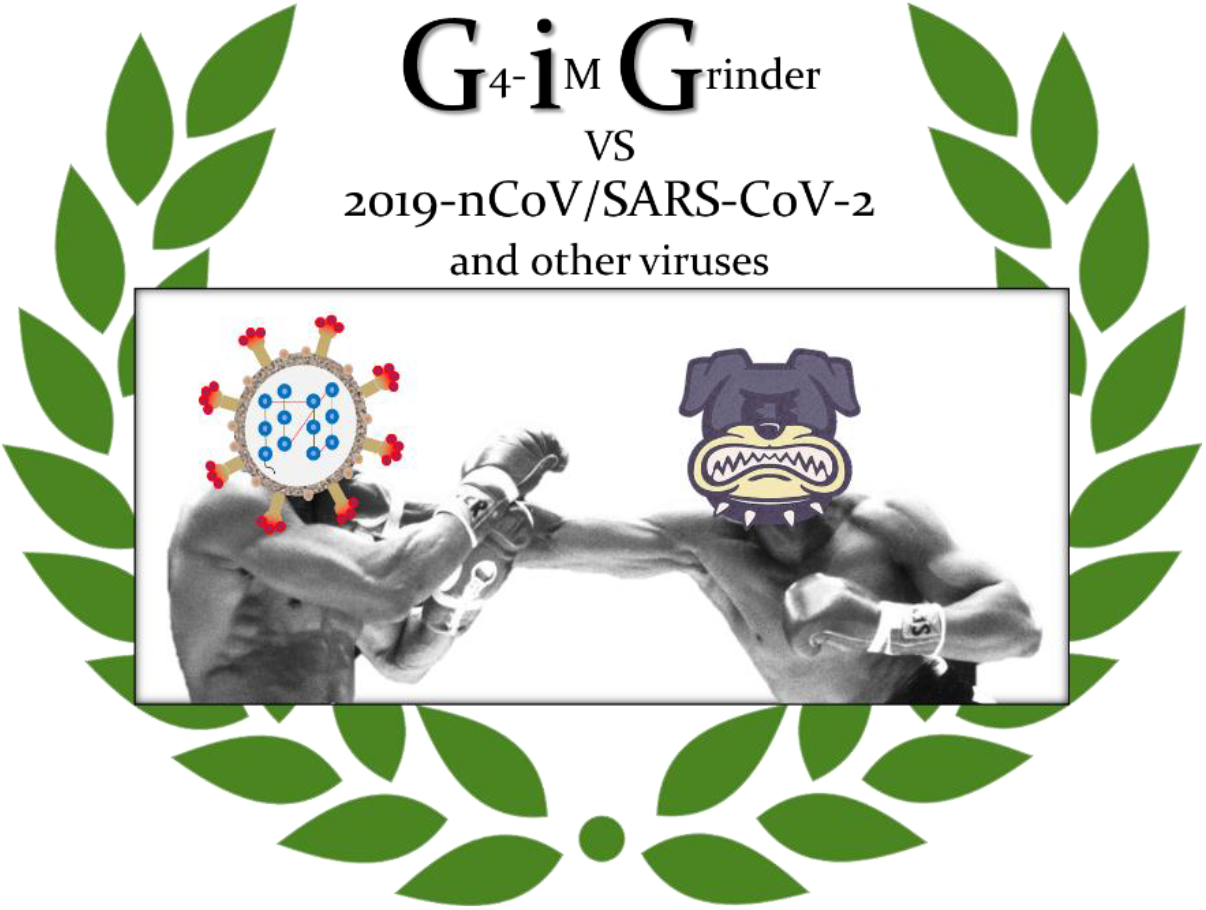

## INTRODUCTION

The severe acute respiratory syndrome coronavirus 2 (SARS-CoV-2 or 2019-nCoV) is a positive-sense single-stranded RNA virus from the *Betacoronavirus* genus, within the *Coronaviridae* family of the *Nidovirales* order. Although it is believed to have originated from a bat-borne coronavirus, (1–3) the 2019-nCoV can spread between humans with no need of other vectors or reservoirs for its transmission. The virus is responsible for the ongoing COVID-19 pandemic that has caused hundreds of thousands of deaths, millions of infected, and a disastrous strain on the economy of most countries and citizens worldwide.

The origin of the virus has been traced back to the Chinese city of Wuhan, where the first cases of infected individuals were reported amongst the workers of the Huanan Seafood Market (4, 5). This wet exotic animal market, where wild animals including bats and pangolins are sold and prepared for consumption, offers ample opportunities for pathogenic bacteria and virus to adapt and thrive. Such circumstances led Cheng and colleagues to predict the current pandemic back in 2007 (6). In their own words: “*the presence of a large reservoir of SARS-CoV-like viruses in horseshoe bats, together with the culture of eating exotic mammals in southern China, is a time bomb. The possibility of the re-emergence of SARS and other novel viruses from animals or laboratories and therefore the need for preparedness should not be ignored*”.

The fight against the 2019-nCoV has now become a global problem. In this current scenario, the scientific community is playing a fundamental role in defeating the virus and minimizing the number of victims. Their work includes, to name a few, the development of fast and reliable detection methods, the identification of therapeutic targets within the virus, and the development of active drugs and vaccines to cure and to prevent infections, respectively.

G-Quadruplexes (G4s) and i-Motifs (iMs) have been proposed as therapeutic targets in many disease aetiologies. G4s are Guanine (G) rich DNA or ARN nucleic acid sequences where successive Gs stack in a planar fashion via Hoogstein bonds to form four-stranded structures, stabilized by monovalent cations (7). iMs on the contrary, are Cytosine (C)-rich regions that fold into tetrameric structures of stranded duplexes (8–10). These are sustained by hydrogen bonds between the intercalated nucleotide base pairs ***C·C***^+^ when under acidic physiological conditions.

The importance of these genomic secondary structures has been abundantly studied during the last years (11–16). They have been found to be regulatory elements in the human genome implicated in key functions such as telomere maintenance and genome transcription regulation, replication and repair (17). G4 structures have also been identified in fungi (18–21), bacteria (22–26) and parasites (27–32). Their occurrence are known in many viruses that afflict humans as well. These include the HIV-1 (33–35), Epstein-Barr (36, 37), human and manatee papilloma (38, 39), herpes simplex 1 (40, 41), Hepatitis B (42), Ebola (43) and Zika (44) viruses. Here they can regulate the viral replication, recombination and virulence (28, 45, 46).

iMs have been less studied in general, especially outside of the human context. With regards to viruses, Ruggiero *et al*. recently published the formation of an iM in HIV-1 (47), whilst we reported the presence of the known cMyb.S (48) iM within the Epstein-Barr virus (49). Despite the lack off reports, iMs present great potential as viable targets against viruses. For example, the *in silico* analysis of the rubella virus revealed an extremely dense genome of potential iMs (density as counts per genomic length) that surpassed its human counterpart by over an order of magnitude (49). Other viruses, such as the measles and hepacivirus C, were also very rich in potential iMs with densities similar to the human genome.

In this work, we wished to contribute to the ongoing effort against the COVID-19 pandemic by investigating the relationship between the 2019-nCoV and quadruplex targets. With this aim, we analysed the prevalence, distribution and relationships of Potential G4 Sequences (PQS) and Potential iM Sequences (PiMS) in its genome. These PQS and PiMS have been assessed according to their potential to form, uniqueness, frequency of appearance, conservation rates between 3297 different 2019-nCoV clinical cases, confirmed quadruplex-forming sequence presence and localization within the genome. The study of the 2019-nCoV and its quadruplex results were expanded to integrate the *Coronaviridae* family, Group IV of the Baltimore classification and the entire virus realm, as to allow a wider range of interpretation. With all this information at hand, our final objective was to identify biologically important PQS and PiMS candidates in the virus. To substantiate our bioinformatic analysis, we analysed experimentally some of these sequences by CD and NMR spectroscopies. Our *in vitro* results confirmed the formation of stable quadruplexes that can form in the viral genome, suggesting that they may be suitable targets for new therapeutic or diagnostic agents (46, 50). Hence, our analysis into the virus realm, and especially the 2019-nCoV, may provide useful insights into using quadruplex structures as targets in future anti-viral treatments.

## MATERIALS AND METHODS

### G4-iM Grinder and G4-iM Grinder’s database upgrade

In this work, we have used the G4-iM Grinder (GiG) package for the analysis of all viruses (49). GiG is an R-based algorithm that locates, quantifies and qualifies PQS, PiMS and their potential higher-order versions in RNA and DNA genomes. In order to extract more information and better analyse the viruses, we first upgraded GiG. Two new functions were developed and are now incorporated in the GiG-package (as of GiG version 1.6.0) named *GiG.Seq.Analysis* and *GiG.df.GenomicFeatures* (<u>supplementary material</u>, section 1). To help us locate any G4 or iM already studied in the literature, we updated the GiG’s DataBase (*GiG.DB*) to version 2.5.1. The library now includes 2851 quadruplex-related sequences that can be identified within any of GiG’s results. The database is categorized by the capability of the sequence to form or not form quadruplex structures (2141 do, 710 do not), their relation to G-based or C-based quadruplexes (2568 G4, 283 iM) and the genome type (1858 DNA, 993 RNA). The reference information (including DOI and/or PubMedID) of each sequence is also listed and accessible to facilitate further studies.

### Pre-analysis. G4-iM Grinder’ parameter configuration

With these upgrades at hand, we retrieved the 2019-nCoV’s reference sequence (GCF_009858895.2) from the NCBI database (51). We also downloaded those of 18 other viruses which can cause mortal illness in humans, including six other pathogenic Coronavirus, as comparison (<u>supplementary material</u>, section 2).

As a workflow, we applied the functions *GiG.Seq.Analysis* (to study their G- and C-run characteristics), *G4iMGrinder* (to locate quadruplex candidates) and *G4.ListAnalysis* (to compare quadruplex results between genomes) from the G4-iM Grinder package (GiG) to all the viruses. The ‘size restricted overlapping search and frequency count’ method (Method 2, M2A and M2B) was used to locate all the potential candidates. Then, these PQS and PiMS were evaluated by their frequency of appearance in the corresponding genome, the presence of known-to-form quadruplex structures sequences within and their probability of quadruplex-formation score (as the mean of G4Hunter (52) and the adaptation of the PQSfinder algorithm (53)). To compare between virus species, we calculated the density of potential quadruplex sequences per 100000 nucleotides 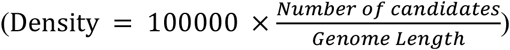.

We previously saw that viruses have a wider-range of PQS and PiMS densities than that of the human, fungi, bacteria and parasite genomes (49). Some were totally void whilst others were very rich in potential candidates. So, we explored different quadruplex definitions to determine the most useful configurations for the analysis of the viruses at hand. These different definitions control the characteristics of what the algorithm considers a quadruplex. They include the acceptable size of G- or C-repetitions to be considered a run, the acceptable amount of bulges within these runs, the acceptable loop sizes between runs, the acceptable number of runs to constitute a PQS or PiMS, and the total acceptable length of the sequence (Figure 1, A). A flexible configuration of quadruplex definitions will detect larger amounts of candidates at the expense of requiring more computing power and accepting sequences that are more ambiguous in forming quadruplex structures *in vitro* (as determined by their score; with longer loops, smaller runs, more bulges and more complementary G/C %, Figure 1, B). More constrained definitions result in the opposite. Hence, for the analysis, we chose three different configurations: a *Lax* configuration (which accepts run bulges and longer ranges of runs, loops and total sizes), the *Predefined* configuration of the package (which restricts sizes but still accepts run bulges), and the original *Folding Rule* (54, 55) (which restricts length and does not accept run bulges) (Figure 1, C Left). Then, we calculated the PQS and PiMS densities of each virus to allow a direct size-independent comparison between them all (Figure 1, D), and filtered the results by their *in vitro* probability of formation score. The score filters were set to |20| and |40| to allow us the study of both the medium (PQS Score ≥ 20; PiMS Score ≤ −20) and the high probability candidates (PQS Score ≥ 40; PiMS Score ≤ −40; Figure 1, B) within the results.

**Figure 1.**
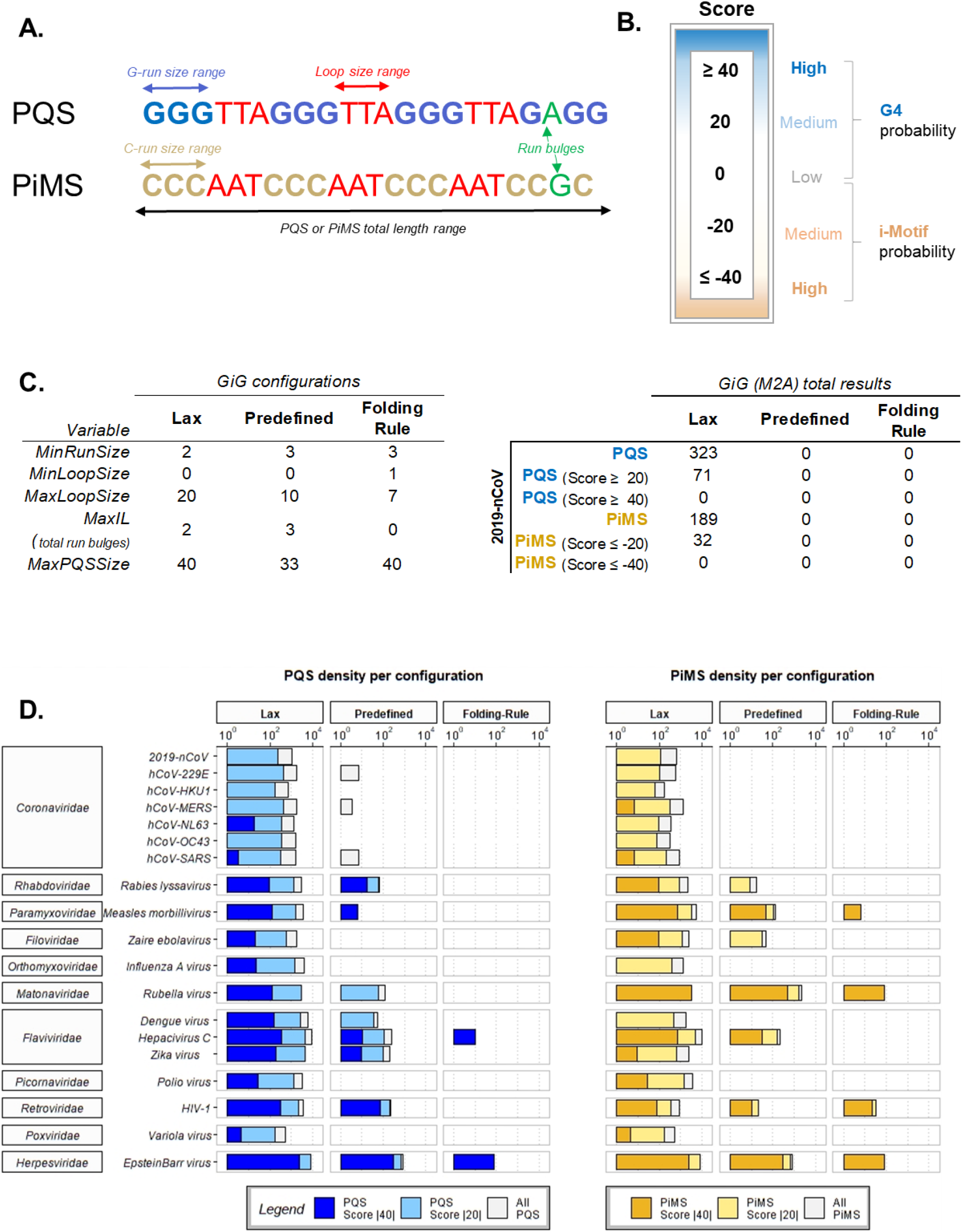
**A**. Results with G4-iM Grinder depend on the quadruplex definitions introduced to the algorithm. Sizes of G- or C-runs, loops and the entire sequence, together with an acceptable number of bulges within the runs are part of the definitions. **B**. The structures found with GiG under the definitions proposed by the user can be evaluated for their *in vitro* probability of formation. This scale is the mean of G4Hunter (that considers G richness and C skewness for PQS or *vice versa* for PiMS as factors) (52) and an adaptation of PQSfinder (that considers run, loop and bulge effects on the structure) (53). Positive values mean that the sequence is more capable of forming G4s, whilst more negative values mean that it is more capable of forming i-Motifs. Values near zero are not good candidates. **C. Left**, GiG’s quadruplex definitions used in this work. Granting more freedom to the quadruplex search will increase the number of structures found, at the expense of requiring more computational power and potentially finding more sequences with ambiguous quadruplex formation potential. **C. Right,** Total results found within the 2019-nCoV by configuration and score criteria. **D.** PQS and PiMS densities (per 100000 nucleotides) found per different configuration and score criteria for 19 viruses that cause mortal illness in humans. X scale is in logarithmic scale (base 10). Results are categorized by their score: intense colours (blue for PQS, yellow for PiMS) are the most probable to form *in vitro* (score over |40|), lighter bars are the density of structures with at least a score of |20| and grey bars are the densities without the score filter (hence accepting all potential structures).

The best configuration to obtain a sufficient amount of results for almost all the viruses analysed was the *Lax* set-up. This was also relevant for the reference genome of the 2019-nCoV (Figure 1, C Right). Given the small size of the viral genomes, the increase in computational power was deemed acceptable and hence, we established this *Lax* configuration as the default configuration for all posterior searches with GiG.

### Analysis with G4-iM Grinder and other tools

The 3297 different 2019-nCoV viral genomes sequenced during the pandemic (from December-2019 to July-2020, by different laboratories worldwide) were retrieved from the online database GISAID (56, 57). These genomes are the result of filtering the database by their coverage (<1 % N content), completeness (>29000 nucleotides) and association to a clinical patient history (only those that have it). All other viral genomes used were retrieved from the NCBI database.

To analyse these genomes, we employed the workflow described in the pre-analysis section using the *Lax* parameter configuration. We investigated the biological features potentially affected by candidates using the function *GiG.df.GenomicFeatures*. The conservation of each PQS and PiMS found in the reference genome was calculated as {Conservation (%) = 100 × *Ng*^+^/∑ *Ng*} where *Ng* is the total number of genomes, and *Ng*^+^ is the number of genomes with the PQS or PiMS candidate. The genomic pairwise alignments, used to study the similarity between viruses and detect PQS and PiMS variations between species, were done using the package *Biostrings* from the Bioconductor repository. To compare potential quadruplex presence and prevalence between genomic groupings (species, families, groups and the entire virus realm), we calculated the genomic density of several arguments. These were calculated using the *GiGList.Analysis* function of the GiG package (density per 100000 nucleotides). The arguments were the density of results (PQS and PiMS), density of results with score filters (with at least |20| or |40|), density of already confirmed sequences that form G4 or iM within, and uniqueness (as {Uniqueness (%) = 100 × *Ns*^f=1^/∑ *Ns*} where *Ns* is the number of sequences, and *Ns*^f=1^ is the number of sequences with a frequency of appearance of 1 in its respective genome). For the G- and C-runs density analysis of the viruses, we used the function *GiG.Seq.Analysis* from the GiG package. The arguments here were: densities of runs with different sizes (two or three to five long G- or C-runs) and with different bulges per run (zero and/or one). Relationships between arguments were calculated through Spearman’s rank correlation (ρ), Pearson correlation (corr) and Kendall’s τ (<u>Supplementary material</u>, section 3, Figures 6 – 10).

### Candidate selection

PQS and PiMS candidates were selected according to their potential to form, uniqueness, frequency of appearance, conservation between 3297 different 2019-nCoV clinical case genomes, confirmed quadruplex presence and localization within the genome.

### NMR experiments

Oligonucleotides (0.3 mM) for NMR experiments were purchased from IDT, and suspended in 200 μl of H_2_O/D_2_O 9:1 in 25 mM KH_2_PO_4_ and 25 mM KCl buffer. Spectra were acquired on Bruker Avance spectrometers operating at 600 MHz, and processed with Topspin software. Experiments were carried out at temperatures ranging from 5.1 to 45 °C and pH from 5 to 7. NOESY spectra in H_2_O were acquired with a 150 ms mixing time. Water suppression was achieved by including a WATERGATE module in the pulse sequence prior to acquisition

### Circular Dichroism (CD)

Circular dichroism (CD) studies were performed on a JASCO J-810 spectropolarimeter using a 1 mm path length cuvette. Spectra were recorded in a 320–220 nm range at a scan rate of 100 nm min^-1^ and a response time of 4.0 s with four acquisitions recorded for each spectrum. Data were smoothed using the means-movement function within the JASCO graphing software. Melting transitions were recorded by the monitoring the decrease of the CD signal at 264 nm. Heating rates were 30 °C/h. Transitions were evaluated using a nonlinear least squares fit assuming a two-state model with sloping pre- and post-transitional baselines. Oligonucleotide solutions for CD measurements were prepared at the same buffer conditions as the NMR experiments. Oligonucleotide concentration was of 50 μM.

## RESULTS

### 2019-nCoV and quadruplexes

G4-iM Grinder’s analysis of the 2019-nCoV reference genome revealed 323 PQS and 189 PiMS candidates. As none scored over |40| (high probability of formation), we focussed on the candidates with scores higher than |20| (medium probability). This filter resulted in the detection of 71 PQSs, 7 of which also scored over 30 (22 and 2 %, respectively; Figure 2, A). Regarding PiMS, 35 scored −20 or less, and 10 below −30 (19 and 5 %, respectively). In both cases, all the candidates were unique (only occuring once in the 2019-nCoV genome) and none included confirmed G4s or iMs (listed in the *GiG.DB*) within their sequence.

**Figure 2.**
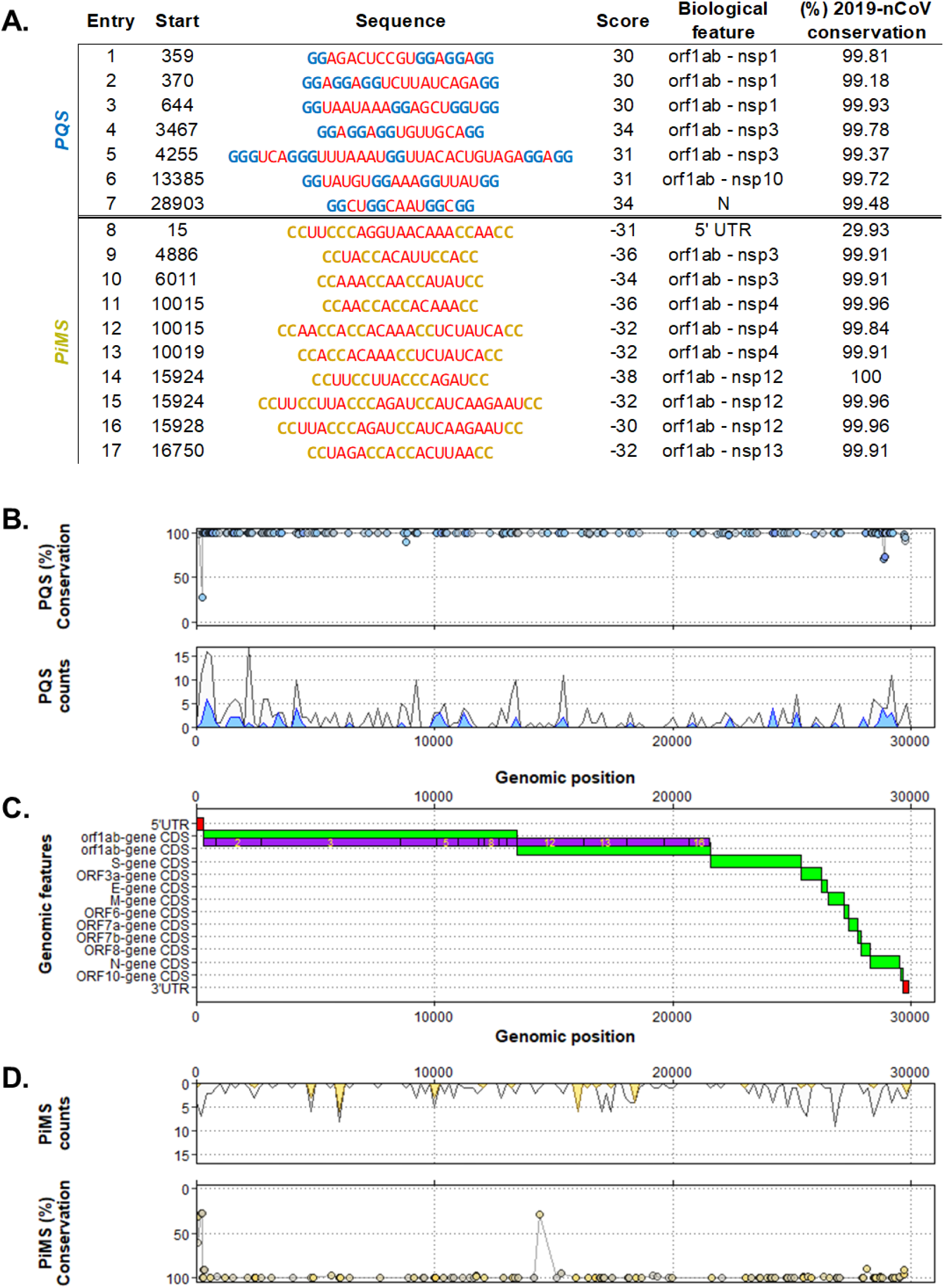
**A**. Top scoring PQS (Score ≥ 30, entry 1 to 7) and PiMS (Score ≤ −30, entry 8 to 17) found in the 2019-nCoV ordered by their localization in the genome. In blue and in dark yellow, the G- and C-runs respectively, and in red, the loops. For each entry, the biological feature column lists the genomic landmark that hosts the potential quadruplex. The percentage of conservation is also given. **B, Top**. PQS count density plot related to the genome position. Y axis is counts per 200 nucleotides (Binwidth = 200). Grey coloured density plots are all the results found, whilst blue density plots are the results found with at least a score |20|. **B, Bottom**. Percentage of conservation of each PQS found along the genome of the 2019-nCoV. Each point represents one PQS. The PQS score is given by the fill colour of the points, where lower scores are greyer, and bluer points have higher (absolute) scores. **C.** Distribution of the biological features of the 2019-nCoV virus by its genomic position. In red, UTR regions, in green, CDS and genes regions, and in purple, nps of the orfab gene. **D, Top**. Percentage of conservation of each PiMS found along the genome of the 2019-nCoV. Each point represents one PiMS. The PiMS score is given by the fill colour of the points, where lower scores are greyer, and higher (absolute) scores are more yellow. **D, Bottom**. PiMS count density plot related to the genome position. Y axis is counts per 200 nucleotides (Binwidth = 200). Grey coloured density plots are all the results found, whilst yellow density plots are the results found with at least a score |20|.

We studied the potential biological effect of these candidates by determining their location and distribution within the genome of the 2019-nCoV. All PQSs with scores over |30|, except one (found in the N gene; entry 7, Figure 2, A), were located in the orf1ab polyprotein gene (in the nsp 1, 3 and 10 areas). Similarly, all the high-scoring PiMS were located in the orf1ab gene (in the nsp3, 4, 12 and 13 areas), except a candidate located in the 5’ UTR region (entry 8, Figure 2, A). When we lowered the score filter to |20|, most results were identified as part of the orf1ab polyprotein. However, some PQSs were also found as part of the S, orf3a, M, orf8 and N genes and some PiMS as part of the S, orf3a, N genes and 3’ UTR. When no score-filter was applied, nearly all biological features presented quadruplex candidates, with the exception of orf7a, orf7b and orf10, as well as the orf6 gene in the case of PIMS (Figure 2, B and D).

For each PQS and PiMS detected on the reference 2019-nCoV, we studied its conservation rates using different 2019-nCoV genomes sequenced by different research groups/laboratories/institutions at different times and locations of the pandemic. In total, we analysed 3297 different 2019-nCoV genomes associated with clinical cases of CoVID with morbid or mortal outcomes. On average, the PQS conservation rate for all candidates was 98.6 ± 7.4 %, whilst for PiMS it was 96.7 ± 12.7 %. For results that score at least |20|, PQS conservation decreased to 98.4 % whilst PiMS increased to 97.4 %.

The relationship between the candidate’s conservation rate and the biological features potentially affected by the PQS and PiMS were then explored. On the one hand, we found that the less conserved sequences are located mainly in the 5’ UTR section, where sequence conservation can fall to 27 %. On the other hand, the 3’ UTR region, showed higher conservation levels with minima of 90 %. The rest of the sequences found in most of the other landmarks presented very high conservation rates, ranging between 99 and 100 % (297 PQS out of 323, 153 PiMS out of 189). Some exceptions exist, however. For example, three PQSs were located in the N gene with conservation rates of 70 – 74 %, and another candidate was found in the orf1ab gene (position 8775, nsp4 region) with a conservation rate of 89 %. In addition, one PiMS found in the orfab gene (position 14391, nsp12 within the Coronavirus RPol N-terminus) was located also with a very poor conservation (29 %).

In a wider context, the total number of PQSs found in the 3297 different 2019-nCoV genomes ranged from 228 to 468 sequences (reference genome is 323). None of them presented already confirmed G4 sequences and all were unique (not repeated in the genome). The total number of results for PiMS were between 133 to 241 sequences (reference genome is 189). Several 2019-nCoV genomes (Hangzhou/ZJU-01-2020, Hangzhou/ZJU-03-2020 and Hangzhou/ZJU-08-2020) presented long repetitions of C-tracks in the start or end regions. In its DNA version, these C-tracks can form the C15, C18, C21 (58) and ATXN2L (15) iMs when protonated. The RNA version of the iM can also form, although with an lower stability (59).

### 2019-nCoV, the *Coronaviridae* family and quadruplexes

We then explored the relationship of the 2019-nCoV with the rest of *Coronaviridae* family members. The most similar genomes found to that of the 2019-nCoV were the SARS-coronavirus (SARS-CoV) and the Bat coronavirus BM48-31/BGR/2008 (Bat-CoV-BM), with pair-alignment score values of 79 and 75 % respectively. These two genomes also presented several PQSs and PiMSs in common with 2019-nCoV. The common candidates were analysed and their potential biological influence within the 2019-nCoV genome determined.

For PQSs, nine sequences are common between all three genomes (2019-nCoV, Bat-CoV-BM and the SARS-CoV), and can be found in the 5’ UTR, 3’ UTR and in the middle of the polyprotein orf1ab regions of the 2019-nCoV. Eleven other sequences are common only to the Bat-CoV-BM and are located within the 5’ UTR region, near to the orf1ab starting positions or in the middle of the orf1ab polyprotein. Eight additional PQSs are common only to the SARS-CoV and appear in the orf1ab gene and 3’ UTR region of the 2019-nCoV.

For PiMS, only one sequence common to all three genomes is located in the middle of the orf1ab gene. Five other PiMSs common only to the Bat-CoV-BM are found in the 5’ UTR region, near to the orf1ab starting positions or in the middle of the orf1ab polyprotein. Nine additional sequences are common only to the SARS-CoV and can be found in the start (less than 40 nucleotides away from the UTR; 2/9) and in middle of the polyprotein orf1ab gene (4/9), the E gene (1/9), and the N gene (2/9) of the reference 2019-nCoV.

The common PQS and PiMS’s position difference between the 2019-nCoV and the SARS-CoV is 59 ± 47 nucleotides, whilst the difference between the 2019-nCoV and the Bat-CoV-BM is 133 ± 109 nucleotides. The scores of these common sequences are however rather poor. None of the PiMS scored less than −20 and only four PQSs scored more than 20 (at least medium probability of formation; Figure 3, A). Of these, the best candidate scored over 30, (entry 6, Figure 2 and entry 2 in Figure 3). In general, the sequences presented small run sizes, a high number of bulges, long loop lengths and a high number of complementary nucleotides within, which lowered their probability for quadruplex-formation.

**Figure 3.**
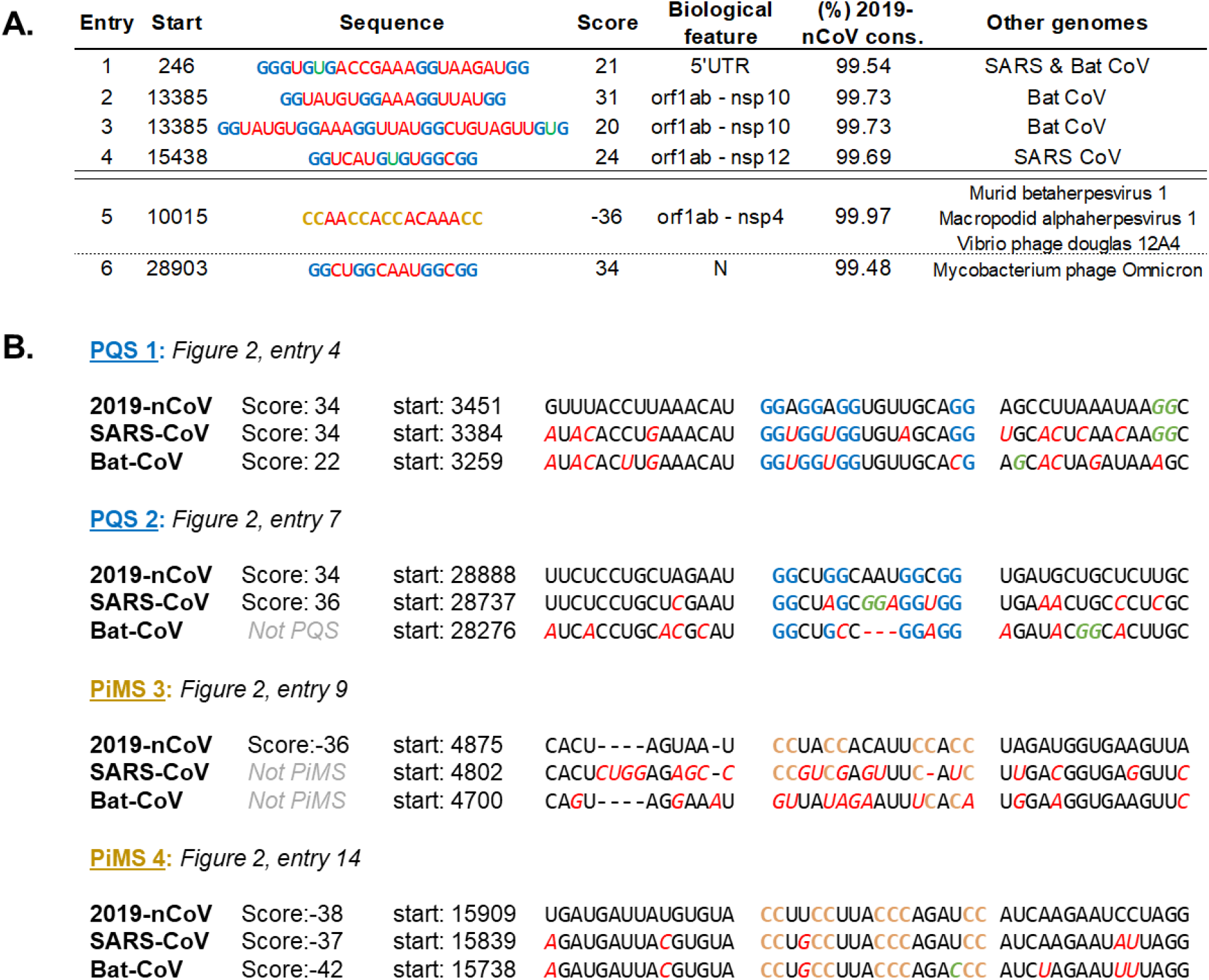
**A.** PQS sequences found with a score of over 20 and common to the 2019-nCoV, SARS-CoV and/or Bat-CoV BM48-31/BGR/2008 (entry 1 to 4), or which are common to other non-related viruses (entry 5 and 6). In blue, the G-runs and in yellow, the C-runs. In red, the loops and, in green, the bulges within the runs. For each entry, the biological feature column lists the genomic landmark that hosts the potential quadruplex in the 2019-nCoV. The percentage of conservation of each entry (between 3297 different 2019-nCoV genomes sequenced at different places and times during the 2019-2020 epidemic) is also given. Conservation is abbreviated Cons. **B**. Alignment of four PQS and PiMS with the highest (absolute) scores found in the 2019-nCoV but not in the SARS or Bat coronavirus. Other common PQS and PiMS were used as fixed points to align the intermediate genomes, with which then to find the locations and variations of these potential structures within all three genomes. Nucleotides in blue and yellow are the potential G- and C-runs respectively that may give rise to the G4 or iMs. In red, the nucleotides that are different from the reference 2019-nCoV genome. In green, G or C nucleotides that may contribute to a nearby sequence.

We focused on the highest scoring sequences found in 2019-nCoV but not common to the other SARS-CoV and Bat-CoV-BM, and identified the variations in their sequences (Figure 3, B). To do so, the previously found common PQS and PiMS were used as fixed points of a genomic-alignment to retrieve the tracks of interest that contain the potential quadruplex sequences.

For PQS1, the changes between the three species occurred in four different nucleotides, three of which are loops. In the Bat-CoV-BM version, the fourth G-run is annulled by the substitution of a G for a C, although a different G-run exists that can still allow the potential quadruplex formation. This PQS, however, has more bulges, longer loops and an increased number of C, which will make it less stable if formed. The 2019-nCoV and SARS-CoV versions have a potential fifth G-run domain downstream which can potentially interact with the PQS.

For PQS2, modifications are greater between species and occur in the central loop and second run area. In SARS-CoV, these modifications allow for a more potentially stable PQS derived from the incorporation of an extra 2-residue G-run. The Bat-CoV-BM version of the PQS has the majority of the central loop deleted and second run invalidated. These modifications, however, do not impede the potential formation of the quadruplex since a nearby G-run downstream can give rise to an alternative PQS.

For PiMS 3, the modifications directly prevent the potential formation of the iM in both SARS-CoV and Bat-CoV-BM genomes. PiMS 4 showed a greater level of conservation. Only two modifications between species occur and are located in the first and third loops. The Bat-CoV-BM version includes an extra C in the last C-run, which makes the PiMS more potentially stable than in the other family versions.

We analysed the potential quadruplex presence and relevance in the family of genomes through a more macroscopic perspective (Figure 4, A). PQS density between *Coronaviridae* species lied between 705 to 3381 PQSs per 100000 nucleotides, with a mean of 1802 (PQS density for 2019-nCoV is 1080). Rousettus bat coronavirus, Rat coronavirus, Parker and Beluga whale coronavirus SW1 had the highest densities in the classification, whilst the human coronavirus HKU1 and several other HKU-CoV had the lowest. All these results were unique and therefore are repeated only once in the genome. 62 % (28 out of 45) of the family presented one or more PQSs with a score of over 40 (most probable PQSs to form quadruplex). The 2G_L1.NAR (27) G4 (in its RNA variant) was found within nine PQSs in the Wigeon-CoV HKU20 genome.

**Figure 4.**
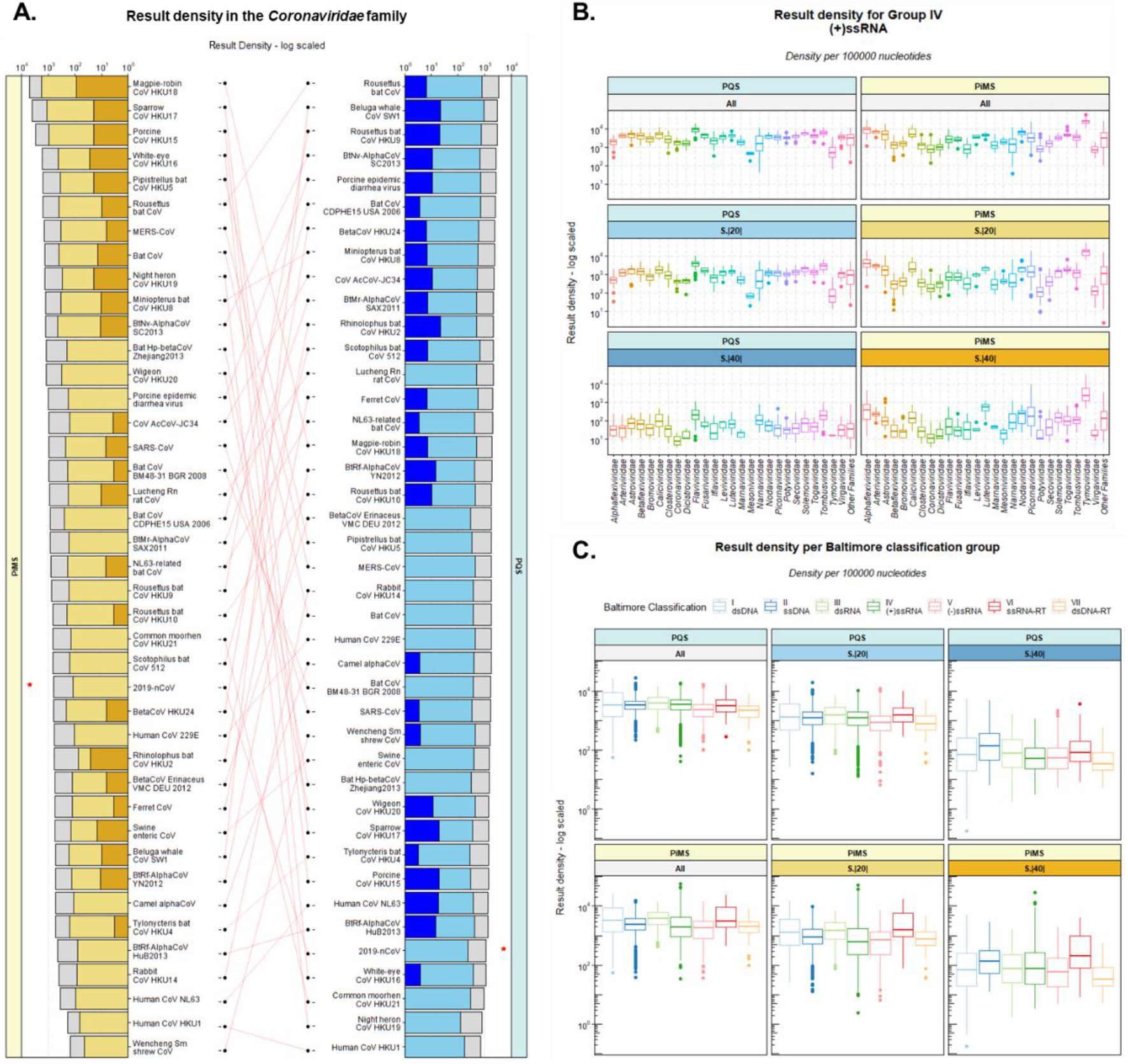
**A.** Result density (per 100000 nucleotides) of PiMS (left) and PQS (right) within the *coronaviridae* family. The results are ordered in descending order of unfiltered result density. In grey, all-result density per virus. In light blue (for PQS) and light yellow (for PiMS), the density of structures with at least medium probability of formation (S.|20|). In intense blue (for PQS) and intense yellow (for PiMS), the density of structures with at least high probability of formation (S.|40|). A base-10 logarithmic scale has been used for the x-axis to appreciate the big differences between results. **B.** Result density boxplots for all the families of viruses in the Baltimore classification group IV (+) ssRNA, which includes the *Coronaviridae* family. All families with less than ten members were merged into the ‘*Other Families*’ group. A base-10 logarithmic scale has been used for the y-axis. Results are divided between PQS (left) and PiMS (right) and between score criteria (top-all result density, middle – result density with a medium probability of formation, bottom-result density with a high probability of formation. **C.** Result density boxplots for all the Baltimore classification groups. A base-10 logarithmic scaled has been used for the Y-axis. Results are divided between PQS (top) and PiMS (bottom) and between score criteria (left-all result density, middle-result density with a medium probability of formation, right-result density with a high probability of formation).

For PiMS, the range for the density of *Coronaviridae* family was in between 142 and 4938 sequences per 100000 nucleotides, with a mean of 1030 ± 929 (2019-nCoV PQS density is 632). Magpie-robin coronavirus HKU18, Sparrow coronavirus HKU17 and Porcine coronavirus HKU15 toped the classification with densities of 4938, 3933 and 2930 respectively. Wencheng Sm shrew coronavirus was the least dense genome, followed by the Human coronavirus HKU1 and Human coronavirus OC43. 64 % of all members (29 out of 45) presented within their results at least one PiMS with a score of −40 or less (most probable PiMS to form *in vitro*). None had already confirmed iMs within their sequence and all were unique, being repeated only once in the genome.

### 2019-nCoV, the Virus Realm and quadruplexes

We further expanded the search to the remaining viruses classified in the NCBI database and found several PQS and PiMS in common with the reference 2019-nCoV. These had a good score and very high *inter*-2019-nCoV conservation rates (Figure 3, A entry 5 and 6). All of these matches were made with Group I viruses (of the Baltimore classification; dsDNA viruses). One PQS candidate located in the N-gene and CDS region of the 2019-nCoV was found in common with a Mycobacteria phage virus. This virus is known to infect the bacterial *Mycobacterium* genus that causes several diseases in humans (tuberculosis and leprosy). The PQS was identified as part of a scaffolding-protein gene within the Mycobacteria phage. Also, a PiMS located in the 2019-nCoV’s orf1ab gene is common to three different viruses; two from the *Herpesviridae* family that infects mammals (marsupials and murids) and a *Podoviridae* virus that infects bacteria from the *Vibrio/Aliivibrio* genus.

In Group IV (which includes the *Coronaviridae* family and 1309 other viruses), we explored the quadruplex presence and distribution per family (Figure 4, B) and calculated their mean and standard deviation (<u>Supplementary material</u>, section 3. Figure 1 to 5). These were then compared to the *Coronaviridae* results.

**Figure 5.**
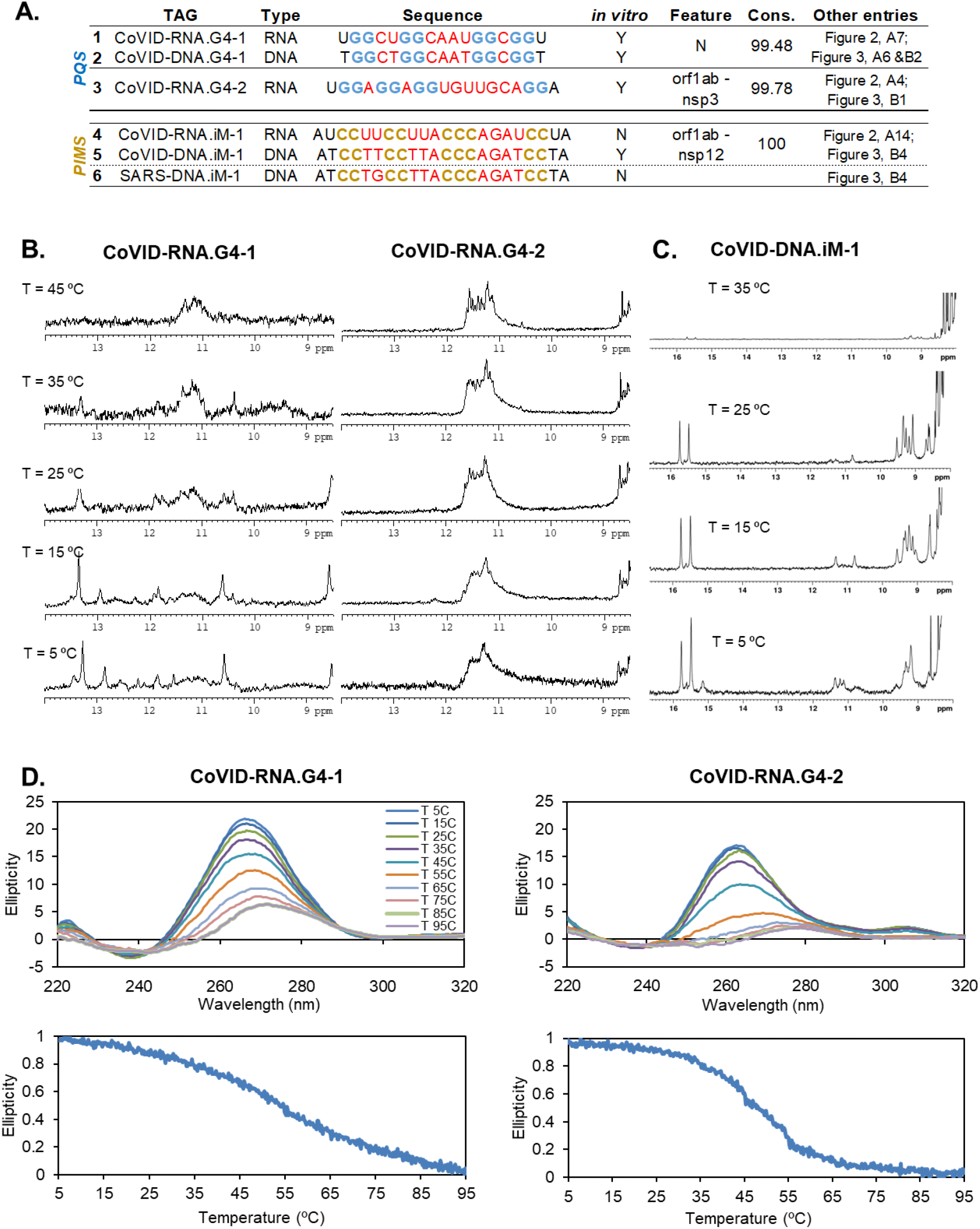
**A**, The candidates examined *in vitro* through biophysical assays. The *in vitro* column states if the sequence forms a quadruplex (Y for Yes, N for No). Feature column is the Biological Feature which hosts the sequence within the 2019-nCoV. Conservation of the sequence found in over 3000 2019-nCoV genomes is abbreviated Cons. **B**, NMR spectra of the two RNA-G4s analysed at different temperatures. **C**, NMR spectra of the DNA-i.Motif analysed at different temperatures. **D**, CD analysis of the two RNA-G4 analysed.

For PQS, the *Coronaviridae* family is in the lower range of the group’s distribution. As a whole, the group’s density was double of that found in the *Coronaviridae* family (means of 4034 and 1802, respectively). To the contrary, the *Flaviviridae* and *Tombusviridae* families showed the highest densities, with genomes that surpassed that of the *Coronaviridae* several fold. In particular for *Flaviviridae*, its mean PQS density was found to be more than four times that of *Coronaviridae* (means of 9079 and 1802, respectively). 106 viruses within the Group also presented already confirmed G4 sequences within their results. These were found in 36 species belonging to the *Flaviviridae* family, 9 to the *Picorbiridae*, 8 to the *Tombusviridae* and 7 to the *Closteroviridae* families, amongst others.

For PiMS, the *Coronaviridae* family was again in the low end of the group’s distribution. The group’s mean density was over three times that of *Coronaviridae*’s (mean of 3731 and 1043, respectively). Contrastingly, the *Tymoviridae* family was found to be very rich in potential candidates (mean of 25359). Its results surpassed by over an order of magnitude most of the other group member results. The *Picornaviridae, Secoviridae* and *Luteoviridae* families also had very dense genomes, and within these four families, 9 virus species presented confirmed iMs in their genome (of 11 in total).

In a wider context, we investigated the prevalence and distribution of potential quadruplexes in the entire virus realm and compared them to Group IV’s results. We also examined the results on a global scale (Figure 4, C).

For PQS, we detected relatively small differences between the groups. Group I (dsDNA) viruses displayed the highest mean density followed by Groups III, II and VI. The entire virus realm density was found within the limits of 40 to 27687 PQSs per 100000 nucleotides with a mean of 4441. The *Herpesviridae* and *Siphoviridae* families from Group I and *Inoviridae* family in Group II displayed the highest PQS densities amongst the 6680 viruses analysed. Additionally, several G4s confirmed in the literature were found in 1372 viruses within Group I, 133 within Group II, 74 within Group III, 106 within Group IV, 18 within Group V, 8 within Group VI and 14 within Group VII (<u>Supplementary material</u>, section 3. Figure 11). Group differences for PiMS were also relatively small, with the largest mean being Group VI followed by Groups I, III and IV. Here, the virus realm density was calculated in the range of 0 to 51771 with a mean of 4221. The *Tymoviridae* family and other Group IV families, mentioned previously, topped the rank, together with some species of the *Herpesviridae* family from Group I. iMs confirmed in the literature were found in 175 viruses within Group I, 5 within Group II, 11 within Group IV, 2 within Group V and 2 within Group VI.

### Biophysical Experiments

To validate the predictions of our bioinformatics search, we selected three candidates using the criteria mentioned in the material and methods section. Two of them were potential G4 forming sequences and the third one was an iM candidate.

The first G4 (CoVID-RNA-G4-1) examined is found in the N-gene of the 2019-nCoV with a conservation rate of 99.48 % (Figure 5, A entry 1). The NMR spectra of this RNA exhibited a broad set of signals around 11 – 12 ppm, characteristic of guanine imino protons involved in G-tetrads. These signals are observed at high temperatures indicating that the G-quadruplex is quite stable (Figure 5, B left). Additional signals around 12 – 13 ppm, which are characteristic of Watson-Crick base-pairs, can also be observed at low temperature. These interactions may arise from loops between G-tracts or alternative conformation such as hairpin-like structures. The CD spectra of the candidate revealed a positive band at 264 nm and negative band with a minimum at 240 nM consistent with the formation of a G4 of a parallel topology (Figure 5, D, left). Melting experiments monitored by CD confirmed the great stability of the G4, whose melting temperature (Tm) was calculated to be 54.4 °C at [K^+^] = 50 mM. Encouraged by these results, we additionally selected another candidate for experimental analysis with a very high conservation rate (CoVID-RNA-G4-2; Figure 5, A, entry 3). This candidate is located in the orf1ab gene within the nsp3 region. As for the previous analysis, NMR and CD spectra revealed a stable parallel G4-quadruplex (Figure 5, B and D, right), with a CD-monitored Tm of 48.1 °C. In this case, the CD spectra presented an additional band at 310 nm, most likely related to the association between two quadruplexes to form a dimeric structure. This is consistent with the number of imino signals observed in the NMR spectra at high temperatures, which suggests the presence of more than two G-tetrads.

In the case of PiMS, we selected a very conserved candidate (100 %) found in the orf1ab – nsp12 region of the virus (CoVID-RNA-iM-1; Figure 5, A entry 4). In NMR, only two small signals appeared in the 12.5 – 14.5 ppm range at neutral pH. Under acidic conditions more signals were observable, including a peak at 15 ppm which could be associated with ***C·C***+ iminos.

However, further analysis by 2D NMR spectroscopy revealed that this signal arises from an AU base pair (<u>Supplementary material</u>, section 4. Figure 13). We must conclude that this sequence, although folded, does not form an iM. This result is not totally unexpected, since the lower stability of RNA vs DNA iM is well known. In spite of this negative result, and to check the capability of our algorithm to detect iMs, we decided to study the DNA version of this sequence (CoVID-DNA-iM-1; Figure 5, A, entry 5). Most interestingly, the NMR spectra of this DNA oligonucleotide exhibited several imino signals in the 15 – 16 ppm range, characteristic of ***C·C***+ base pairs. These signals are observable in the 5.5 to 6.7 pH range (Figure 5, C, and <u>Supplementary material</u>, section 4. Figure 14). Additionally, amino groups from ***C·C***+ (in the 9 – 10 ppm range) and TT base pairs (in the 11 ppm region) are also observable. As TT base pairs are common capping groups in iMs, it is interesting that the SARS version of this sequence (SARS-DNA-iM-1; Figure 5, A, entry 6), which only differs from 2019-nCoV in a single nucleotide within the first loop (from TT to TG), was unable to form an iM under all pH conditions studied (even at pH 5.4).

## DISCUSION AND CONCLUSIONS

### G4-iMGrinder and settings

In this work, we have used G4-iM Grinder to analyse the genome of the 2019-nCoV, and that of many other viruses, in search off potential quadruplex (both G4 and iM) therapeutic targets. To do so, we first expanded G4-iM Grinder’s quadruplex identification and characterization repertoire with two new functions, *GiG.Seq.Analysis* and *GiG.df.GenomicFeatures*. Other functions such as *G4iMGrinder* and *GiGList.Analysis* were upgraded to better analyse and summarise the quadruplex results obtained. Furthermore, over 2800 quadruplex-related sequences were searched for in the literature and included in G4-iM Grinder’s database to rapidly identify confirmed G4s and iMs within all results.

An initial study of the 2019-nCoV genome and 18 other pathogenic viruses revealed the special characteristics that need to be considered for quadruplex-related examinations in these organisms. For most, the original folding rule (which accepts no bulges within the runs and is very constrained in its quadruplex definitions) and the predefined parameters of G4-iM Grinder (which allows more liberty by accepting bulges and longer loops) are too strict to find associated runs that can give rise to quadruplexes. Although other organisms such as *Plasmodium falciparum* or *Entamoeba histolytica* may be less rich in G and C content (49), the size of these genomes enables finding rich G- or C-tracks that can ultimately form potential quadruplexes. In most viruses, however, this does not take place because of the small size of the genomes (in the range of tens to hundreds thousand nucleotides versus the tens of millions for the parasites mentioned, and thousands of millions for humans). Furthermore, most of the G4s found in viruses are complex sequences, with short runs and bulges (for example, HIV-1(33, 35) and Ebola(43)), which elude detection when following traditional quadruplex definitions. To overcome these problems, we took advantage of the great modulability of G4-iM Grinder, and developed, tested and successfully employed a *lax* quadruplex definition configuration for the analysis. With these settings, the number of candidates found increased greatly and included the complex sequences expected in viruses, at the expense of needing more computational power.

### 2019-nCoV

With all these updates and configurations at hand, we focused on the reference 2019-nCoV and located 323 PQS and 189 PiMS unique (only occurring once in the genome) sequences dispersed unevenly in the genome. 20 % of these candidates had at least a medium probability of formation (score over |20|). These were concentrated in the orfab gene (especially nsp 1 and 3 areas for PQS; and nsp 3, 4 and 12 for PiMS), the N-gene and S-gene (a highly variable gene that binds to the ACE2 membrane receptor and controls the viral penetration into the cell (3)). The orf3a (related to virulence by necrotic death inducement and cytokine expression (60)), M-gene (which encodes membrane glycoprotein (61)), orf8 and UTR regions also presented these candidates. Here they may play their biological role if formed. Other genes, such as orb7a and b, and orf10 were found totally void of any quadruplex candidates.

We calculated the 2019-nCoV candidate’s quadruplex conservation rates and quadruplex-related region variability under three different scopes. First, the attention was set exclusively on the virus in an intra-species analysis comprising 3297 genomes of the 2019-nCoV sequenced at different places and times of the pandemic. Here we saw that most of the sequences analysed presented very high conservation rates, although a few candidates in the 5’UTR, orf1ab and N landmarks displayed low conservation with respect to the reference genome (with a minimum of 27 %).

The search was then expanded to the rest of the *Coronaviridae* family. 53 2019-nCoV PQS and PiMS candidates were found in common with the SARS-CoV and/or Bat coronavirus BM48-31/BGR/2008 (Bat-CoV-BM), all of which are suspect of having bats as hosts during their evolution. These common sequences were located in the 3’UTR, N and E genes of the 2019-nCoV, although most were positioned in the orfab gene, and especially in the 5’UTR region. Paradoxically, the candidates found in the 5’UTR site (which regulates the translation of the RNA transcript) include the least conserved group of candidates of the inter-species analysis (with conservation rates as low as 27 %), while also hosting a very conserved family-wise group of candidates (<u>Supplementary material</u>, section 3, Figure 12). One set of sequences are highly conserved within species and family, and another is very un-conserved altogether. On the one hand, high conservation in candidates (maintained through natural selection) may be an important factor for the survival of the virus. This importance may transcend beyond the 2019-nCoV and into other familiar species were PQS and PiMS were found in common. On the other hand, variability in the region may also play a vital role in the ability of the virus to adapt to new hosts, situations and environments.

The highest scoring candidates found in 2019-nCoV were however not common to any other *Coronaviridae* member species. So, we investigated the differences between them through genome alignments and found that most of the sequence versions amongst species (6 out of 8) were still able to form potential quadruplex structures even with modifications. Therefore, these PQS and PiMS, although different from those in the 2019-nCoV, maintain their potential biological role and importance.

Expanding the search for common candidates to the entire virus realm, we matched one PQS and PiMS from the 2019-nCoV with the potential quadruplexes found in four viruses from Group I belonging to the *Herpesviridae, Podoviridae* and *Siphoviridae* families (all dsDNA).

### 2019-nCoV and the virus realm

With G4-iM Grinder, we analysed the entire virus realm in a similar fashion to other studies in the literature (62, 63). However, we used a *lax* definition of quadruplexes to detect G- and C-structures and focused the comparison of the realm to the 2019-nCoV.

Whilst the 2019-nCoV did not present any of the published quadruplex sequences listed in the *GiG.DB* within its genome, other viruses including a wigeon-afflicting Coronavirus did. In the entire virus realm, 1725 viruses presented at least one confirmed G4 sequence in their genome, while 195 at least one confirmed iM sequence (the dimensional discrepancies between both results may partially be due to the difference in the number of G4 and iM entries in the database; 2568 and 283 respectively). The sheer volume of species with confirmed quadruplex structures in all groups of viruses suggests that quadruplexes may be common and necessary genomic regulatory elements for viruses to “live”, thrive and adapt; as seen in other organisms such as humans. However, the prevalence is not homogeneous and varies broadly at the group level although not that much at the family level. For example, some families like Group I’s *Herpesviridae* and *Sphaerolipoviridae*, Groups IV’s *Matonavirirdae* and *Flaviviridae* and Groups II’s *Spiroviridae* presented the highest PQS densities; whilst Groups V’s *Aspiriviridae* and *Fimoviridae*, Groups IV *Mononiviridae* and *Mesoniviridae* and Group’s I *Mimiviridae* displayed the lowest. PiMS showed a similar tendency with Group I (*Sphaerolipoviridae* and *Herpesviridae*) and especially IV (*Tymoviridae*, *Matonaviridae* and *Gammaflexiviridae*) families being the densest in candidates; whilst Groups IV (*Monoviridae* and *Yueviridae*), Groups V (*Fimoviridae* and *Phasmavirirdae*) and Groups I families (*Mimiviridae*) displayed the lowest. These results indicate that viruses/families (and particularly single-stranded ones) are probably more oriented to a kind of quadruplex structure in a group/genome-type independent manner, whilst being contingent upon cation concentration and pH of the environment for formation.

Altogether, the 2019-nCoV displayed general poverty regarding quadruplex candidates when compared in a macroscopic perspective to the virus realm. Its PQS and PiMS density were in the lower end of results from the *Coronaviridae* family, which itself was in the lower end of the (+) ssRNA Group IV (in an approximate ratio of 1:2:4 for PQS and 1:2:8 for PiMS). When put into the entire virus realm context, the 2019-nCoV PQS density was lower than 5813 other viruses analysed (out of 6680), whilst PiMS density was lower than 6125. Nevertheless, other viruses with similar densities to that of the 2019-nCoV were found to possess confirmed G4 and iM sequences within, supporting the potential these structures have for targeting the 2019-nCoV.

### Candidate confirmation *in vitro*

We, therefore, selected the best candidates to evaluate *in vitro*. The highly conserved candidate CoVID-RNA-1 formed a parallel G4 stable even at 45 °C. This G4, located in the N-gene, can possibly interact with the viral RNA packaging, transcription and replication functions of the virus (64). The second sequence, CoVID-RNA-2, also formed a stable parallel quadruplex structure. In this case, the quadruplex monomers interacted amongst themselves to form a higher order structure. CoVID-RNA-2 is located in the nsp3 region of orf1ab very near its SUD domain. This area has been associated with the increased pathogenicity of the virus compared to other *Coronaviridae* that do not present it (65). Additionally, it has been suggested that the SUD domain interacts with G-quadruplexes of the host. These results, however, open the possibility of an intrinsic gene modulation that may be linked with an increased virulence. Such a hypothesis can be extended to the SARS-CoV, as another stable PQS candidate was found in its genome in the same location (Figure 3, B1).

For PiMS, the DNA version of a candidate located in the orf1ab gene of the 2019-nCoV and with a 100 % conservation rate formed an iM at almost neutral pH. However, the SARS-CoV version of the iM (which differs by one nucleotide in the first loop, from TT to TG) was unable to form even at pH 5.1. As TT base pairs are common capping positions, the substitution of the T might prevent the folding in SARS-CoV. Additionally, the presence of C in G4s lowers overall stability of the quadruplex as C can base pair with G and ultimately hinder G-quartet formation (66). For C-based structures, the opposite but with the same effect might also be happening. When we analysed the RNA version of the 2019-nCoV iM, it did not form a quadruplex structure. Despite the fact that the sequences found in 2019-nCoV have an intermediate probability of formation, RNA iMs are known to be less stable than their DNA-versions (59). Still, G4-iM Grinder methodology identified several more candidates with the potential to form iMs in the virus. These results prove that especially for DNA, G4-iM Grinder can be used to find and characterize iMs in even C-poor genomes.

Overall, these results greatly expand the current knowledge we have regarding quadruplexes and the 2019-nCoV (67), and open the door for targeting viruses in general, and the 2019-nCoV in particular, through the use of these nucleic sequences as therapeutic targets in future anti-viral treatments.

## SUPPLEMENTARY MATERIAL

The supplementary material is available online and includes information regarding the genomes used, how to access the results and additional figures.

## ACKNOWLEDGEMENTS

The authors thank Dr. Matilde Arévalo, Rafael Ferreira and Sarah Heselden for their help regarding this topic.

## Authors contributions

Conceptualization, code development, data analysis and visualization, E.B-R; Biophysical assays, I.S and C.G; Draft preparation, E.B-R; Draft review and edition, E.B-R, J.G, C.G. and M.B-L; Funding acquisition: C.G., J.G and M.B-L.

## FUNDING

This work was supported by the 2014-2020 North Portugal Regional Operational Program (NORTE 2020) and the European Regional Development Fund (ERDF) under Grant NORTE-01-0145-FEDER-000019, by the Fundação para a Ciência e a Tecnoloxía (FCT), ERDF and NORTE 2020 through the Grant NORTE-01-0145-FEDER-031142 (Local specific treatment of triple-negative-breast-cancer through externally triggered target-less drug carriers – MagtargetON –), and by the 2014-2020 INTERREG Cooperation Programme Spain–Portugal (POCTEP) through the project 0624_2IQBIONEURO_6_E. C. G. also acknowledges support from the Spanish Ministry of Science, Innovation and Universities (MCIU) under project BFU2017-89707-P.

## CONFLICTS OF INTERESTS

The authors declare no conflict of interest.

## Notes

https://github.com/EfresBR/G4iMGrinder

### Competing Interest Statement

The authors have declared no competing interest.

