## Supplementary Material for "Exploring G and C-quadruplex structures as potential targets against the severe acute respiratory syndrome coronavirus 2"

**Contents table**

1. Methodological scheme and new functions S2
2. Genomes used S4
3. Other bioinformatic figures S5
4. Other biophysical figures S17
5. Data results S19

### 1. Methodological scheme and new functions

Methodological scheme

Proposed methodology used in this work. The genome sequences of the viruses analyzed (including the 2019-nCoV) were analyzed with G4-iM Grinder. G4-iM Grinder detects nucleotides that can form runs and finds the association between runs that can give rise to potential quadruplex sequences (PQS and PiMS). These candidates are then evaluated by the criteria mentioned in the article. On the one hand, these results can then be compared with other viruses through the calculus of the genomic density ($Density = 100000 \times \frac{N^{\underline{o}}\ of\ Results}{Genome\ Length}$). On the other hand, the best candidates can be confirmed *in vitro* through biophysical assays.

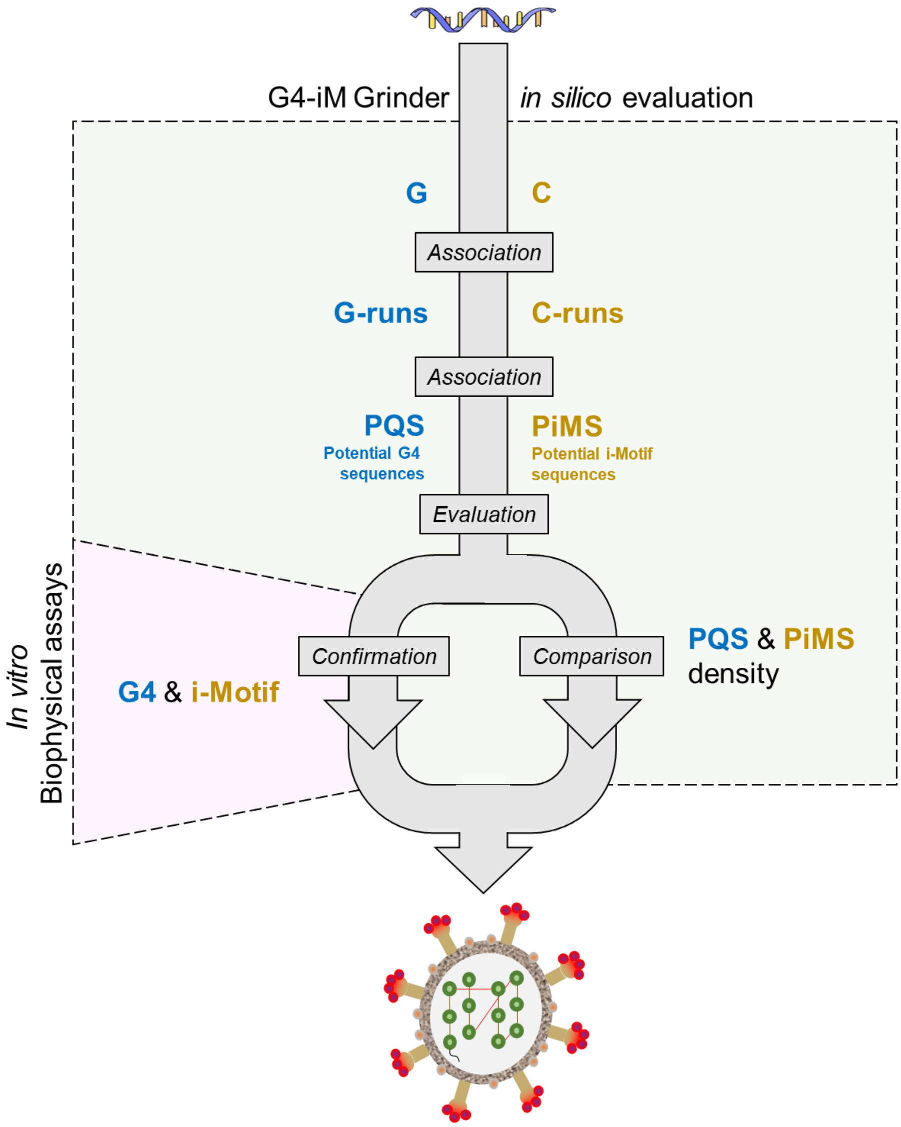

<u>New Functions for G4-iM Grinder</u>

*GiG.Seq.Analysis* function is useful for retrieving the genomic characteristics of the sequence to analyse with GiG. This function can be used before a GiG analysis to determine the best search parameters to obtain quadruplex-related results. The function’s outcome is a data frame with the most relevant genomic features, including length (in nucleotides), type of genome (DNA or ARN), strands (single or double), and G, C, T/U, A and N composition (as % of total sequence). The function also calculates the total number of runs with different conditions (predefined parameters, bulges per run: zero and one-quantities; run lengths: two to five and three to five-length) in the genome, and returns it to the user as total counts or genomic density.

*GiG.df.GenomicFeatures* function is suitable for determining the genomic features that share their location with (and hence may be affected by) GiG’s PQS and PiMS results. It employs the online database connector package “*biomartr*” (51) to retrieve the genomic annotations file for the sequence, with which to then match positions. The function returns a data frame of all the matches found for the input sequences and includes different attributes (IDs, keys, relationships with other features and comments) of the matched genomic features.

### 2. Genomes used

The reference genome of the 2019-nCoV (assembly ascension was GCF_009858895.2 released 13 January 2020) was downloaded from the NCBI database (https://www.ncbi.nlm.nih.gov/). All other genomes used, except those otherwise stated, were downloaded from the NCBI database via the biomartr R package. Further information regarding these genomes can be found within the data results section (section 5).

The 3297 different genomes used in this work of the 2019-nCoV were retrieved from the GISAID database (https://www.gisaid.org/). They were selected by their completeness, their N content, and their association with a clinical case. Further information regarding these genomes can be found within the data results section (section 5).

### 3. Other bioinformatic figures

**Figure 1.** Non-log scaled version of Figure 4 in the manuscript. Potential quadruplex density found in the *Coronaviridae* family (**A**), other Group IV families (**B**), and all the groups analyzed (**C**). Graphs are divided by quadruplex type (PQS and PiMS) and by a score filter (high probability of formation [Score ≥ |40|], medium probability [Score ≥ |20|], and no score filter [All]). Density per 100000 nucleotides.

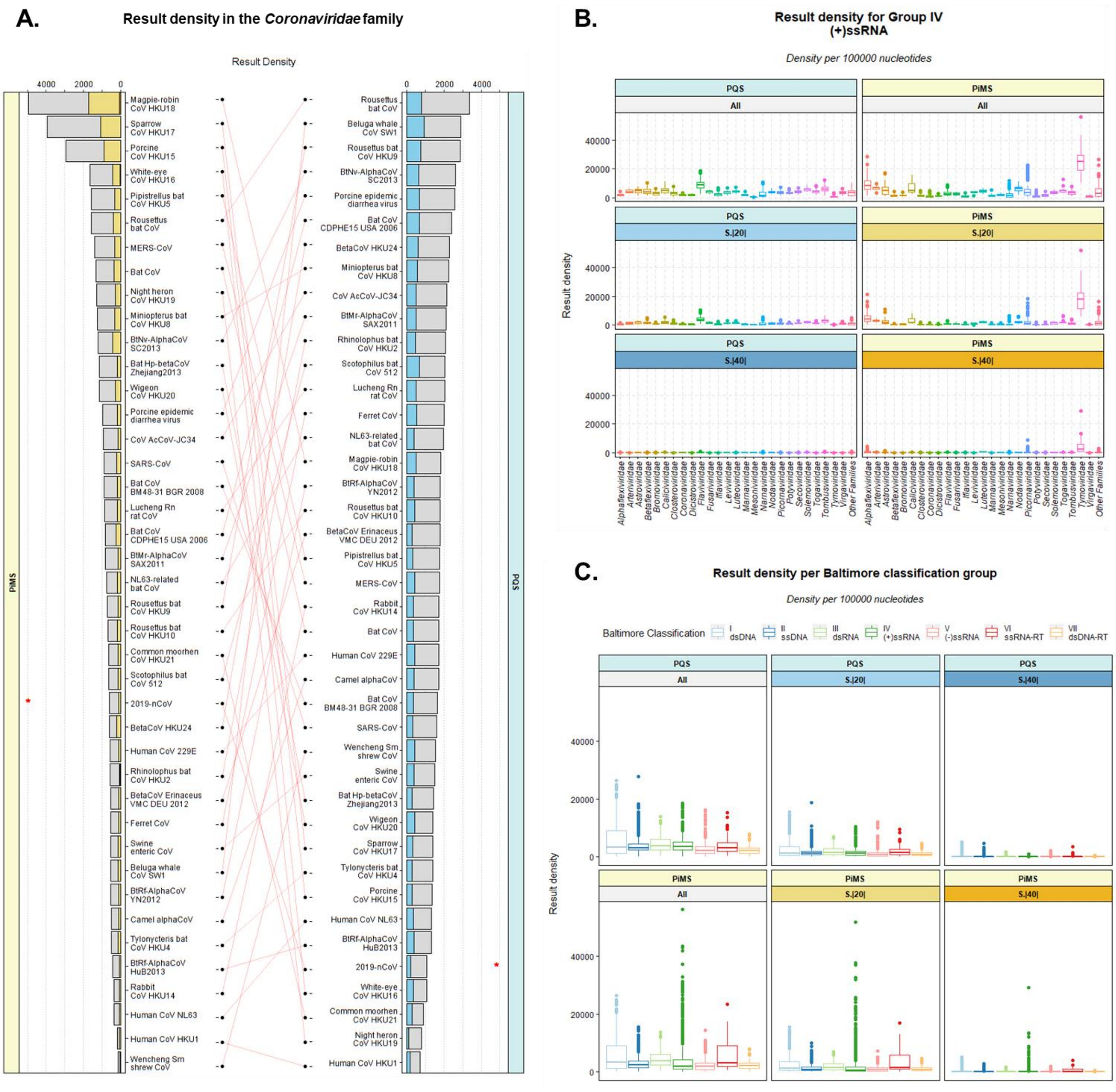

**Figure 2.** Group IV family potential quadruplex density mean ± standard deviation. Graphs are divided by quadruplex type (PQS and PiMS) and by a score filter (high probability of formation [Score ≥ |40|], medium probability [Score ≥ |20|], and no score filter [All]). Density per 100000 nucleotides.

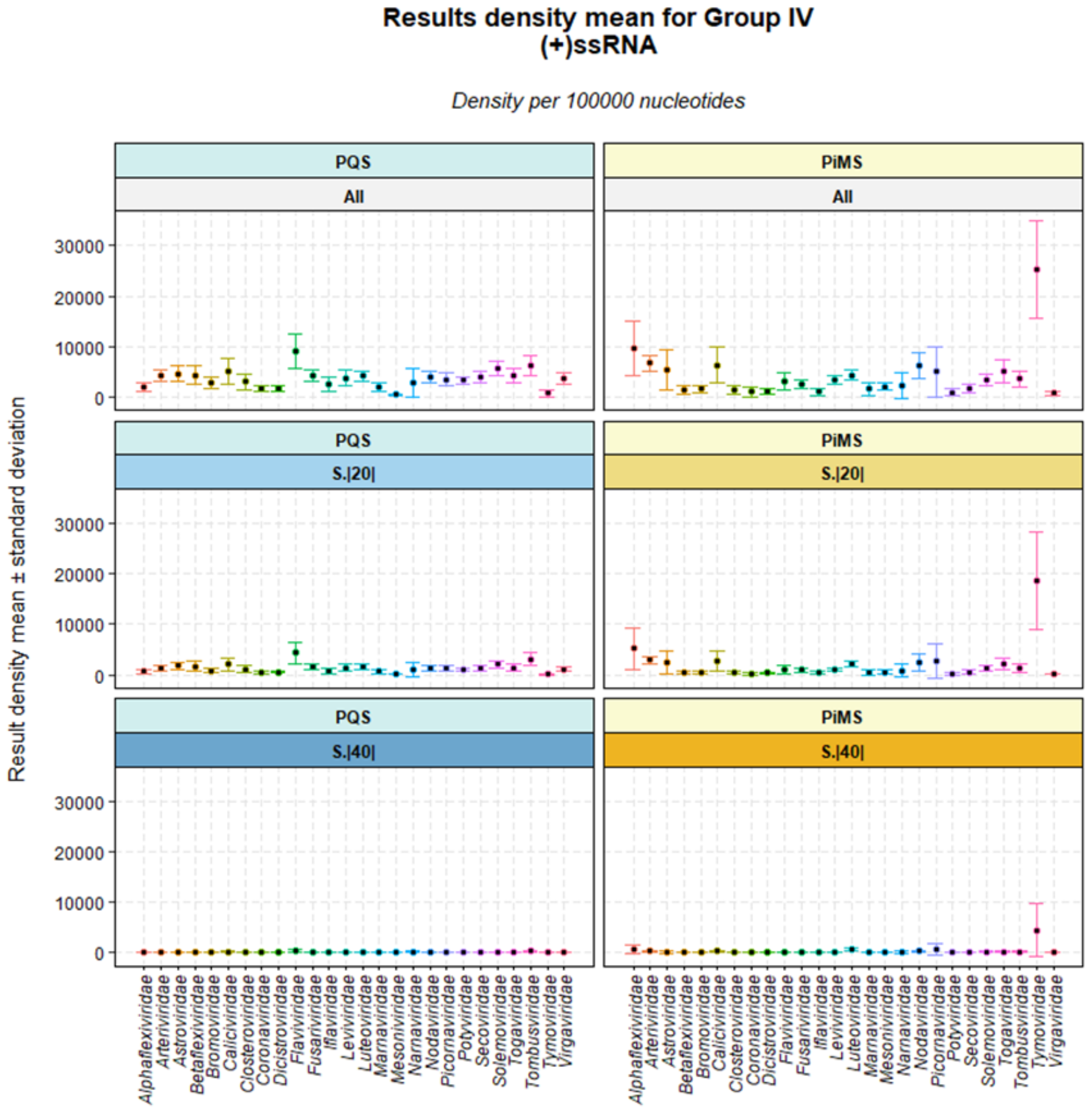

**Figure 3.** Group IV family potential quadruplex density mean ± standard deviation. Y-axis are log scaled. Graphs are divided by quadruplex type (PQS and PiMS) and by a score filter (high probability of formation [Score ≥ |40|], medium probability [Score ≥ |20|], and no score filter [All]). Density per 100000 nucleotides.

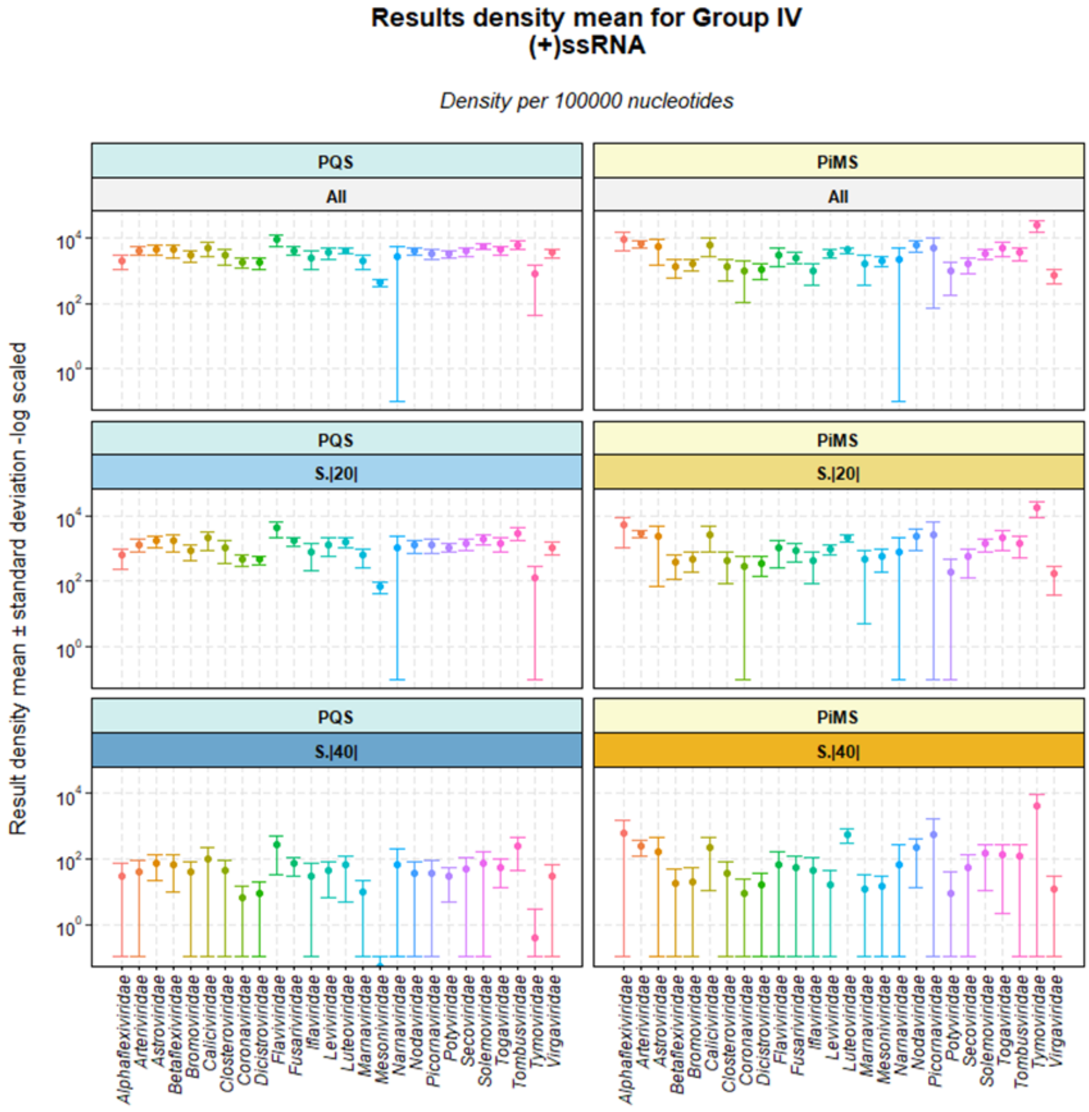

**Figure 4.** All group's potential quadruplex density mean ± standard deviation. Graphs are divided by quadruplex type (PQS and PiMS) and by a score filter (high probability of formation [Score ≥ |40|], medium probability [Score ≥ |20|], and no score filter [All]). Density per 100000 nucleotides.

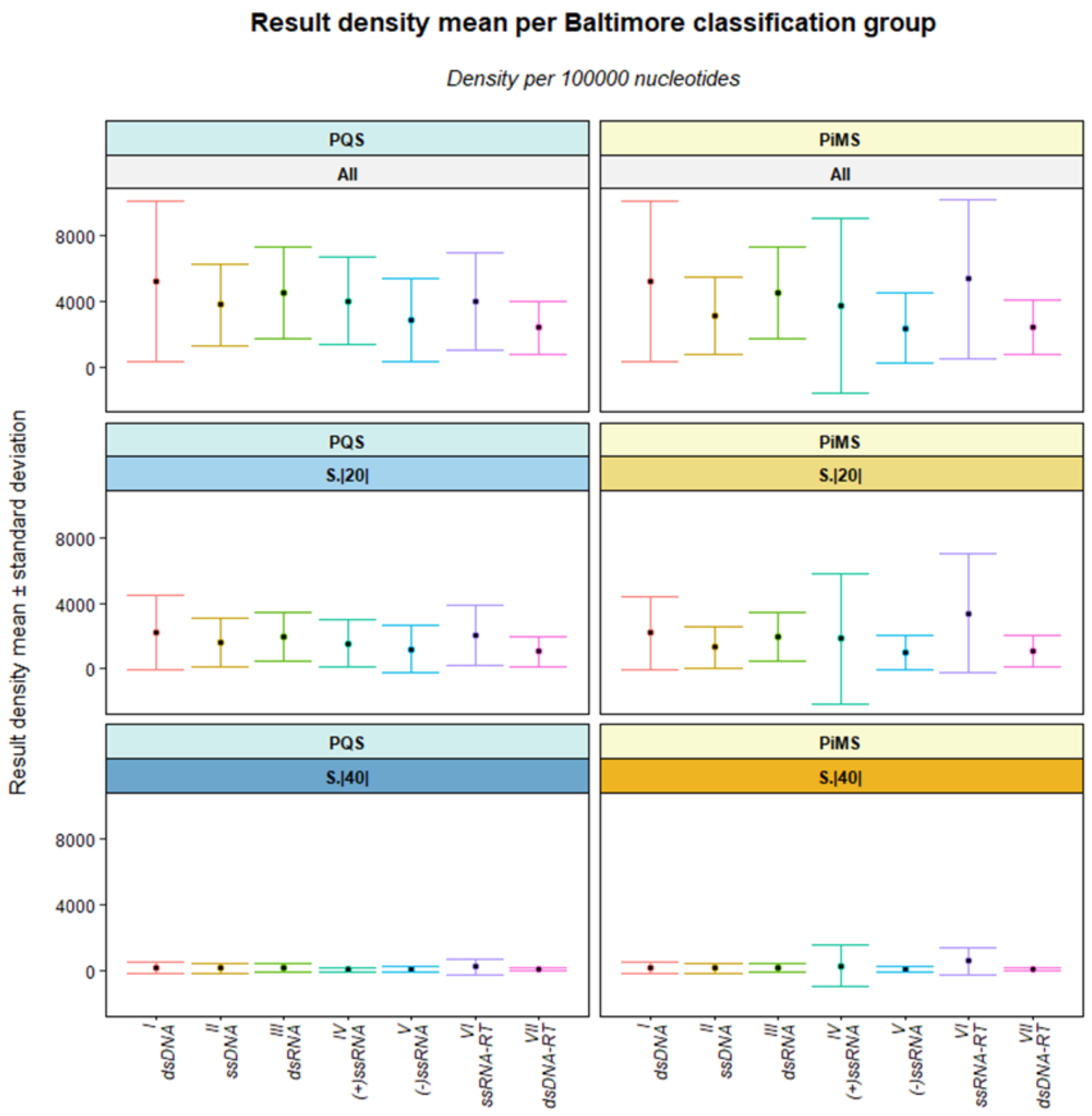

**Figure 5.** All group's potential quadruplex density mean ± standard deviation. Y-axes are log scaled. Graphs are divided by quadruplex type (PQS and PiMS) and by a score filter (high probability of formation [Score ≥ |40|], medium probability [Score ≥ |20|], and no score filter [All]). Density per 100000 nucleotides.

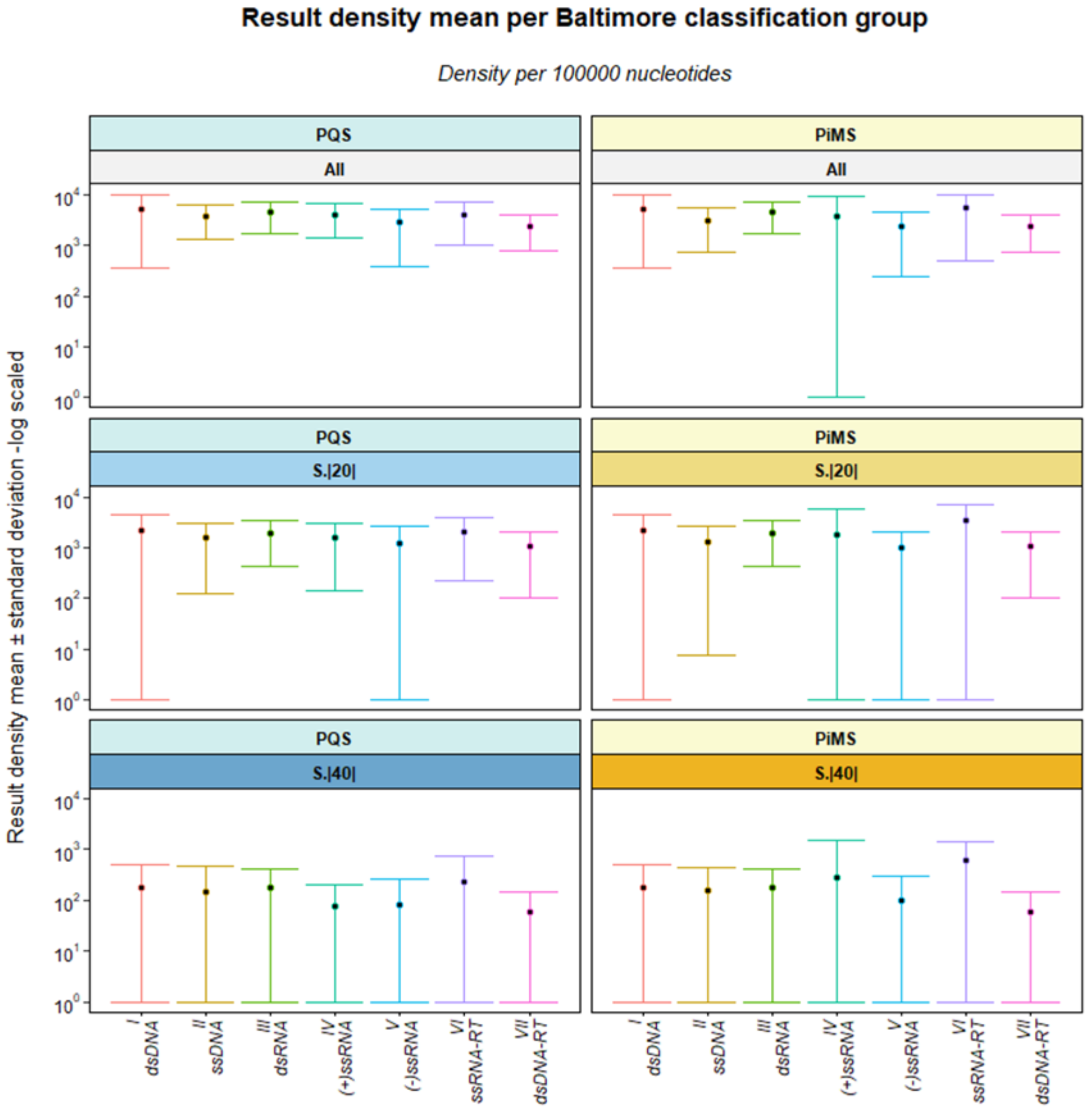

**Figure 6. A.** Potential quadruplex counts (results) versus genome length of the 6680 viral genomes analyzed. **B.** Potential quadruplex density versus G|C % genomic content of the 6680 genomes analyzed. Graphs are divided by quadruplex type (PQS or PiMS) and a score filter (high probability of formation [Score ≥ |40|], medium probability [Score ≥ |20|], and no score filter [All]). For A, axie are log-scaled. Graphs include 2-dimensional density lair and best-fit line. Correlation parameters and their significance are also given. Density per 100000 nucleotides.

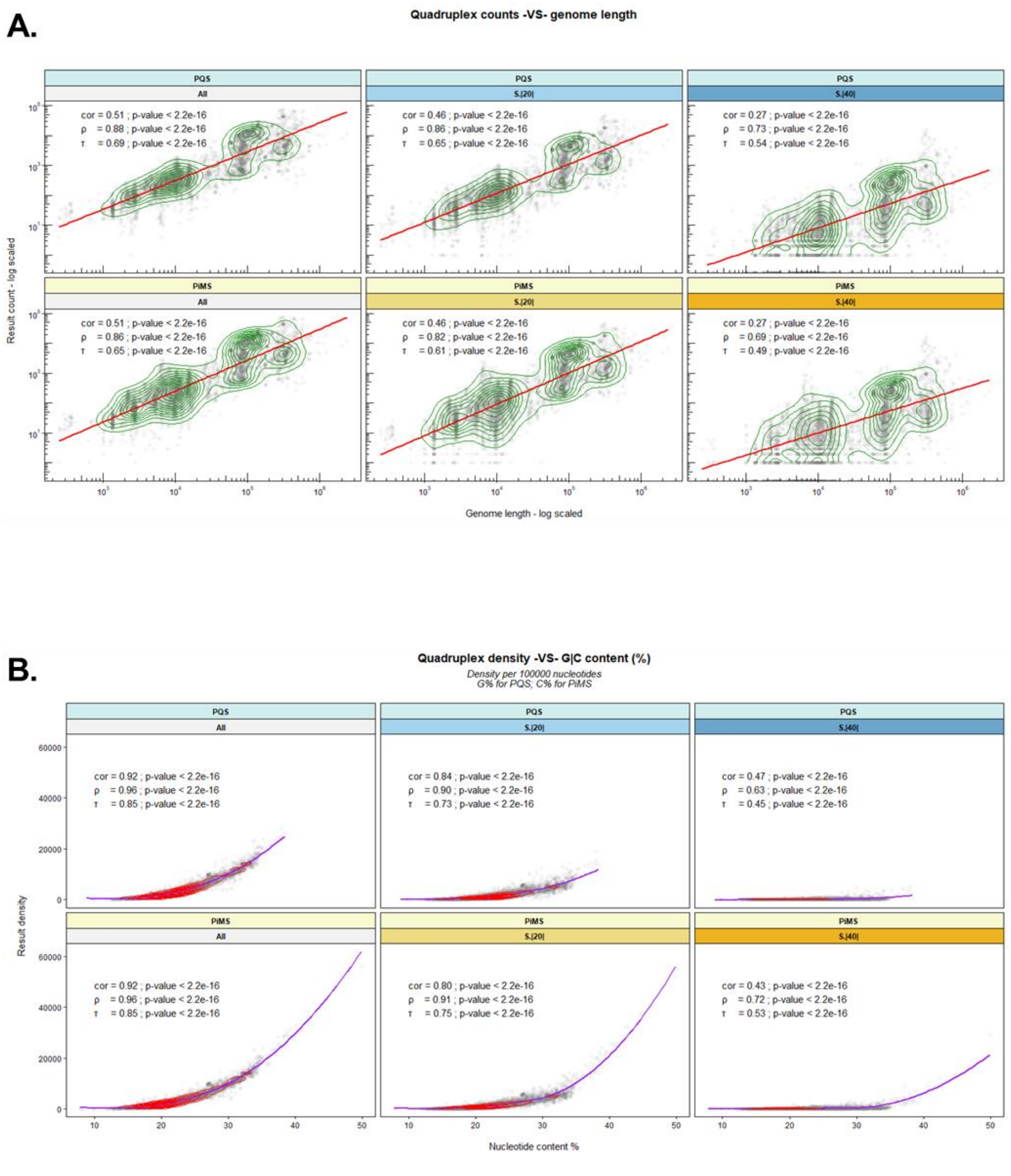

**Figure 7. A.** Run counts versus genome length of the 6680 viral genomes analyzed. **B.** Run density versus G|C % genomic content of the 6680 genomes analyzed.

Graphs are divided by quadruplex type (PQS and PiMS). For B, the graphs are also divided into perfect (in purple - left) and all runs (in green – right). Perfect runs include G and C-runs with no bulges, and with lengths comprehending between two and five nucleotides. All runs include perfect plus imperfect runs (with a bulge per run). For A, axes are log-scaled. Graphs include 2-dimensional density lair and best fit linear model line. Correlation parameters and their significance are also given. Density per 100000 nucleotides.

A.

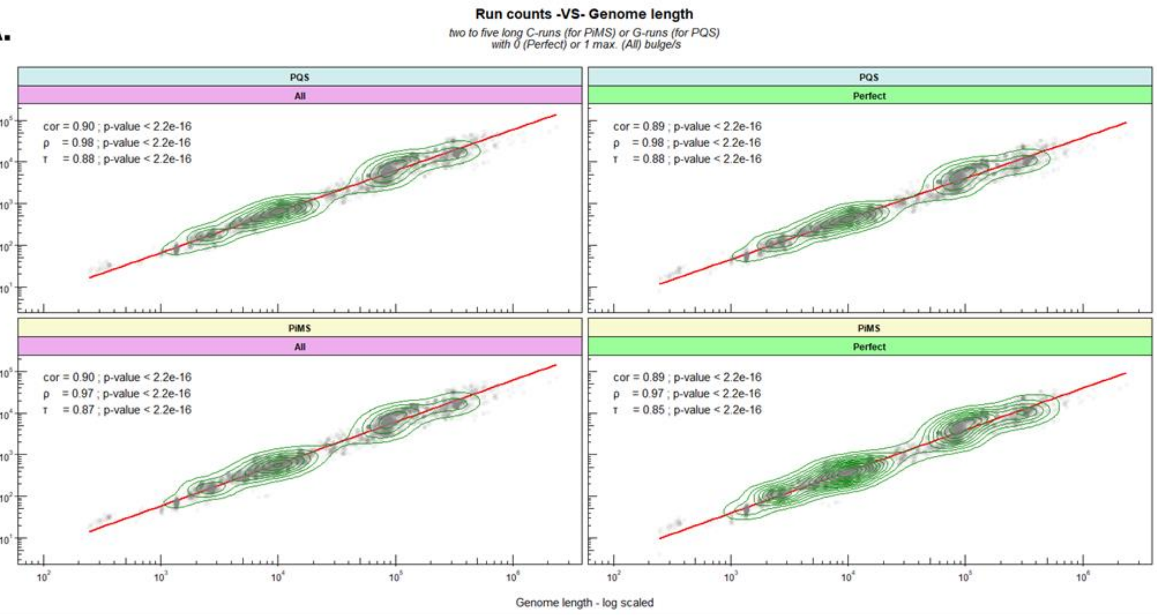


B.

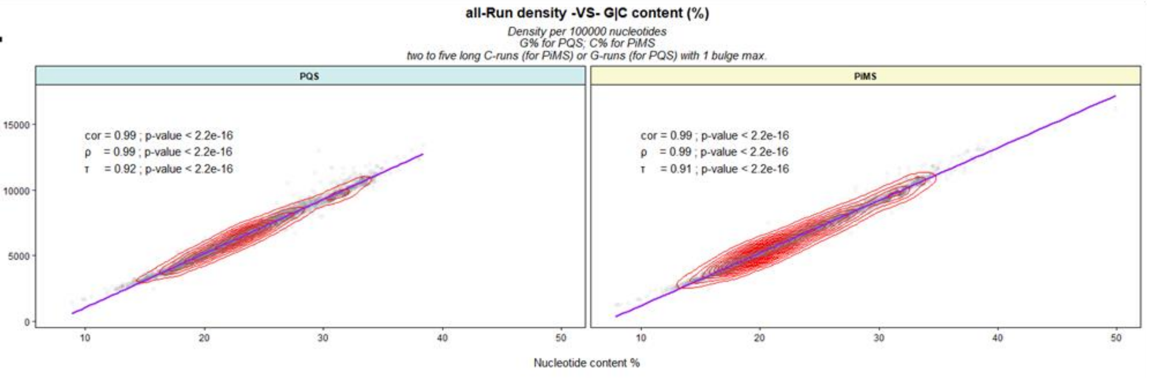

**Figure 8.** Potential quadruplex density of the 6680 viral genomes analyzed versus perfect-run (**A**) and all-run (**B**) density. Perfect runs include runs with no bulges and with lengths between two and five nucleotides. All runs include perfect plus imperfect runs (with a bulge per run). Graphs are divided by quadruplex type (PQS or PiMS) and score (high probability of formation [Score ≥ |40|], medium probability [Score ≥ |20|], and no score filter [All]). Graphs include 2-dimensional density lair in red, and best-fit model line in purple. Correlation parameters and their significance are also given. Density per 100000 nucleotides.

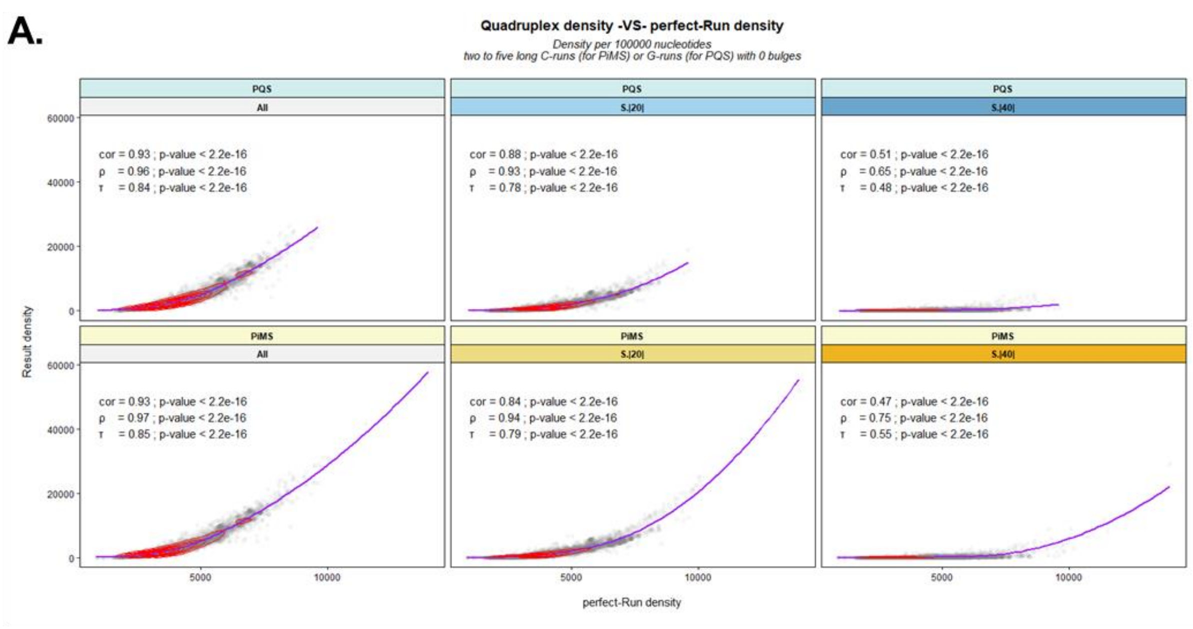


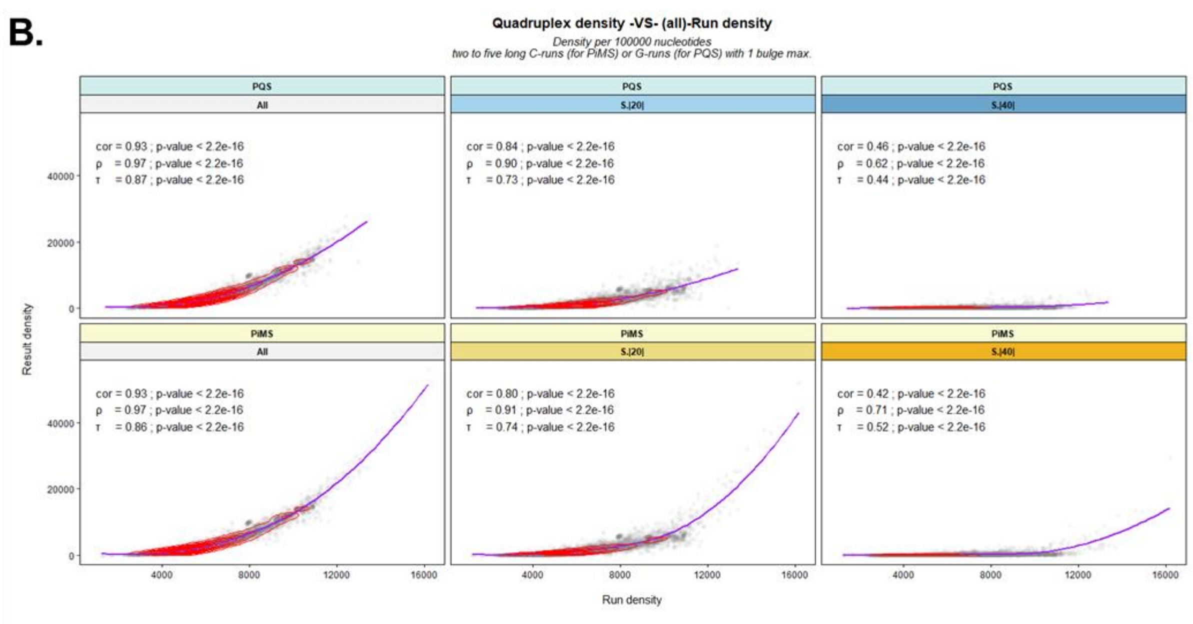

**Figure 9. A.** Potential quadruplex density versus potential quadruplex uniqueness (%) of the 6680 genomes analyzed. **B.** Genome length versus uniqueness (%) of the 6680 genomes analyzed.

Graphs are divided by quadruplex type (PQS and PiMS). For A, the graphs are also divided by a score filter (high probability of formation [Score ≥ |40|], medium probability [Score ≥ |20|], and no score filter [All]). Uniqueness is the number of candidates found in the genome with a frequency of appearance of one, compared to all the candidates found in the genome. Y-axes are log scaled. Graphs include 2-dimensional density lair and best-fit linear model line. Correlation parameters and their significance are also given. Density per 100000 nucleotides.

**A.**

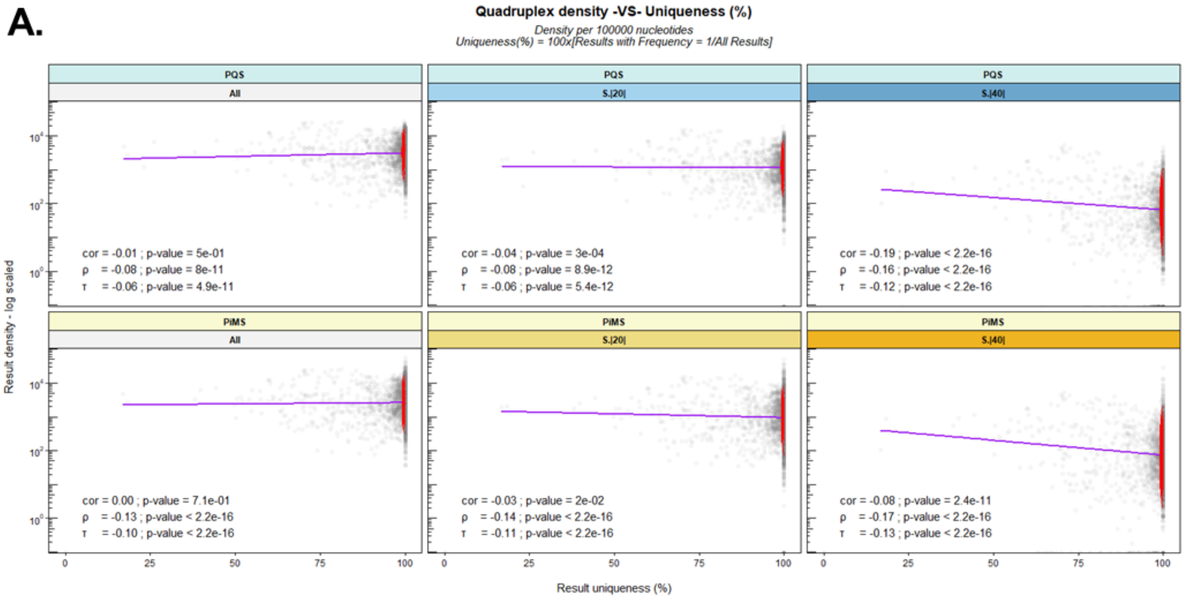


**B.**

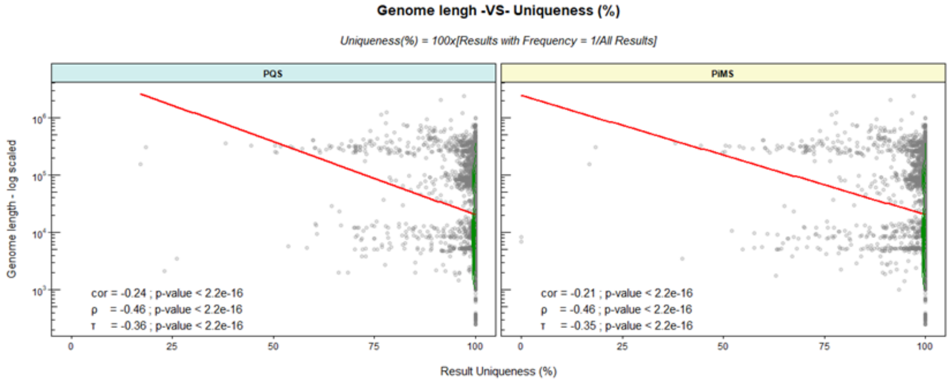

**Figure 10.** PQS versus PiMS densities for all the 6680 viral genomes analyzed. **A.** A global best-fit line is given in purple. **B**. Each group best-fit line is given separately. Graphs are divided by a score filter (high probability of formation [Score ≥ |40|], medium probability [Score ≥ |20|], and no score filter [All]). Axes are log-scaled. Graphs include 2-dimensional density lair. Correlation parameters and their significance are also given. Density per 100000 nucleotides.

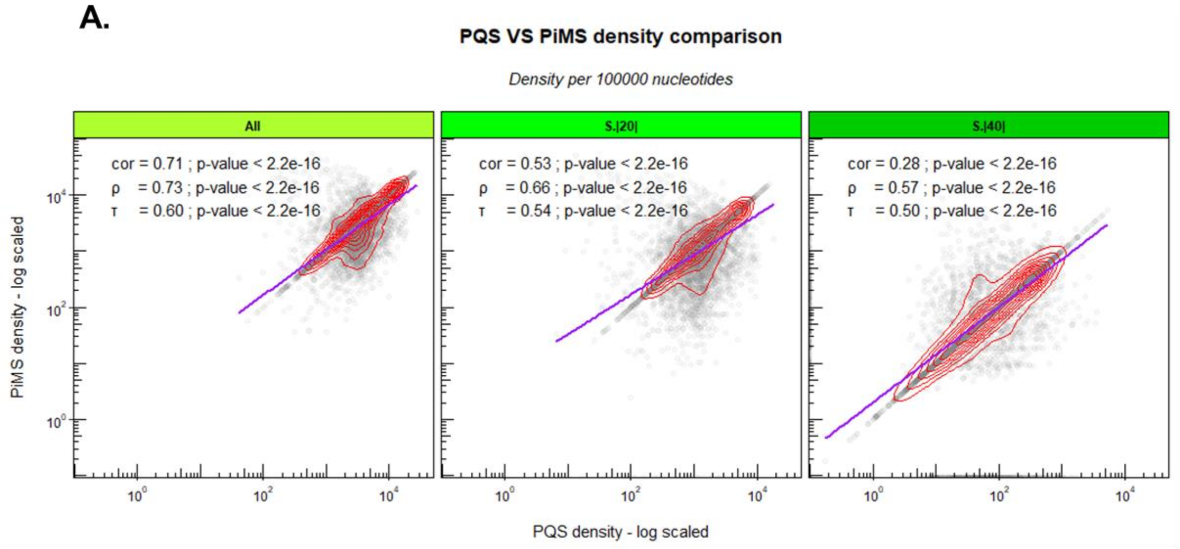


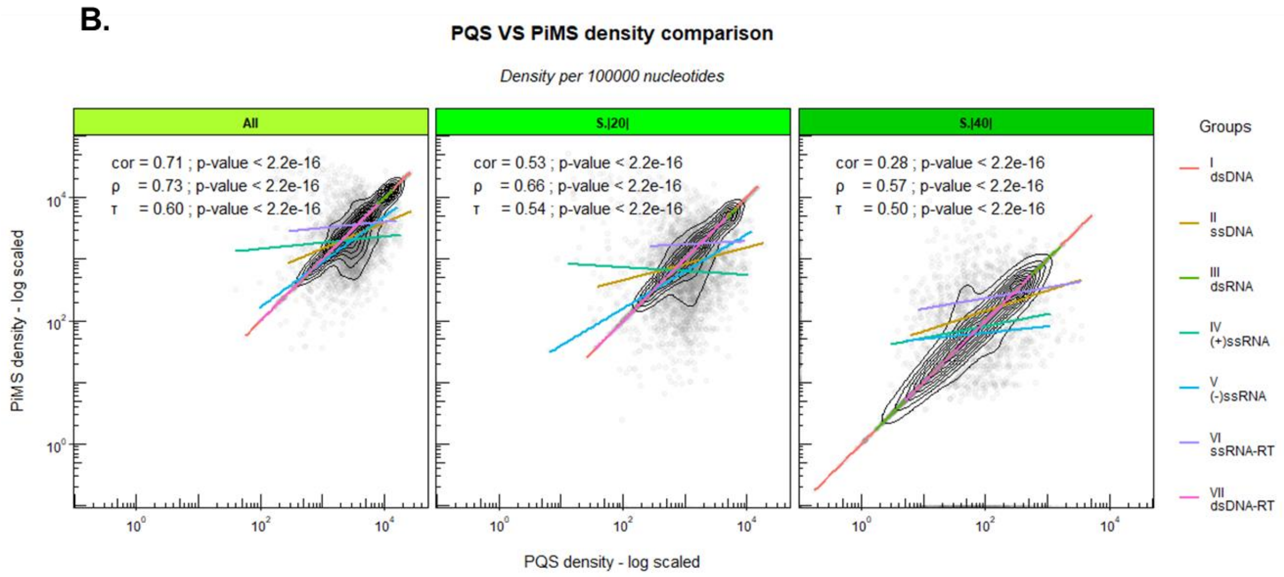

**Figure 11.** Number of species with confirmed G4 and i-Motif forming sequences within (in their DNA and/or RNA versions) per viral group. Graphs are divided by quadruplex type (G4s and i-Motifs). Results are based on *GiG.DB* version 2.5.1.

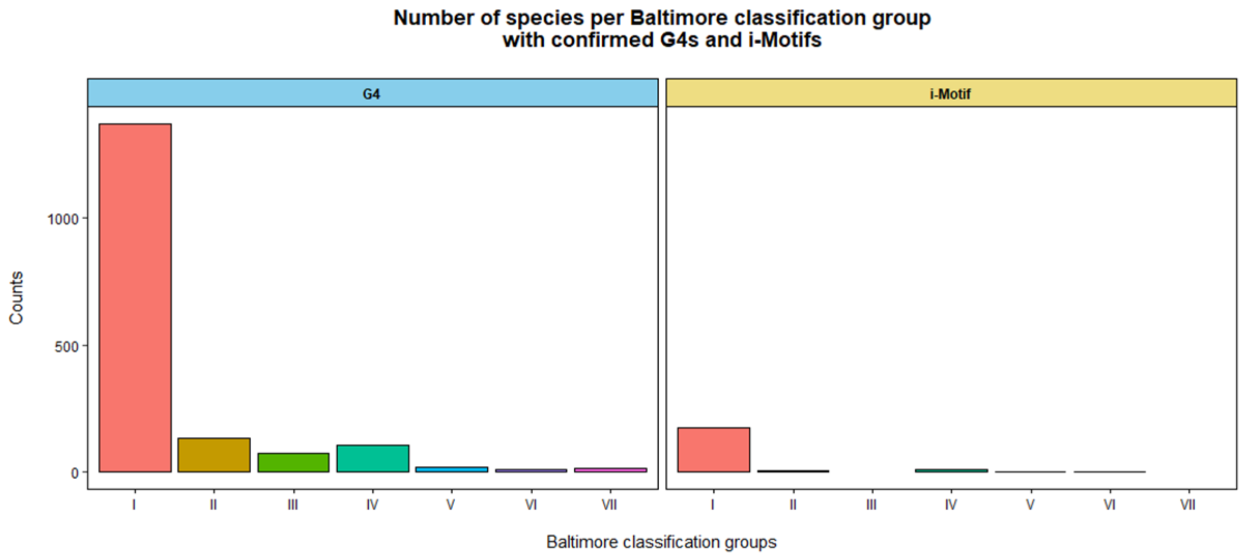

**Figure 12. A.** PQS and **B.** PiMS candidates found in the 5'UTR region of the 2019-nCoV. For both potential quadruplex structures, all candidates are located in two different clusters. For the second clusters, more concatenated candidates exist and continue past the UTR region into the orfab gene. In blue and yellow, G and C-runs respectively. In green, the run bulges. In red the loops. Conservation refers to the degree of conservation (as %) in all the 3297 different 2019-nCoV genomes analyzed. **C.** Some candidate variations between 2019-nCoV of the high scoring but poorly conserved PiMS found in the 5'UTR.

**A.**

```
Cluster 1:                                                                    Score    Conservation
Start    86
         GUGUGGCUGUCACUCGGCUGCAUGCUUAGUG                                      1        97.8

Cluster 2:                                                                    Score    Conservation
Start    236
         GGUUUCGUCCGGGUGUGACCGAAAGG                                           17       27.8 * ^
         GGUUUCGUCCGGGUGUGACCGAAAGGUAAGAUGG                                   18       27.8 * ^
         GGUUUCGUCCGGGUGUGACCGAAAGGUAAGAUGGAGAG                               15       27.8 * ^
                   GGGUGUGACCGAAAGGUAAGAUGG                                   21       99.5 * ^
                   GGGUGUGACCGAAAGGUAAGAUGGAGAG                               16       99.5 * ^
                   GGGUGUGACCGAAAGGUAAGAUGGAGAGCCUUGUCCCUGG                   12       99.4 *
                     GUGACCGAAAGGUAAGAUGGAGAG                                 6        99.5 * ^
                     GUGACCGAAAGGUAAGAUGGAGAGCCUUGUCCCUGG                     4        99.4 *
                           GGUAAGAUGGAGAGCCUUGUCCCUGG                         10       99.5 *
                           GGUAAGAUGGAGAGCCUUGUCCCUGGUUUCAACGAG               4        99.4 *

* common also to Bat-CoV-BM, GCF_000887595.1
^ common also to SARS-CoV GCF_000864885.1
```

**B.**

```
Cluster 1:                                                                    Score    Conservation
Start    15
         CCUUCCCAGGUAACAAACCAACC                                              -31      29.9
         CCUUCCCAGGUAACAAACCAACCAACUUUCGAUCUC                                 -19      29.7
           CCCAGGUAACAAACCAACCAACUUUCGAUCUC                                   -14      31.5
                  CCAACCAACUUUCGAUCUCUUGUAGAUCUGUUCUC                         0        60.7

Cluster 2:                                                                    Score    Conservation
Start    206
         CCGUGUUGCAGCCGAUCAUCAGCACAUCUAGGUUUCGUCC                             -4       27.7
              CCGAUCAUCAGCACAUCUAGGUUUCGUCCGGGUGUGACC                         -2       27.7
                          CCGGGUGUGACCGAAAGGUAAGAUGGAGAGCCUUGUCCC
                                                                              -12      99.4 *

*common also to Bat-CoV-BM, GCF_000887595.1
```

**C.**

TTATA CCUUCCCAGGUAACAAACCAACC AACUU

(Very low conservation: 29.9 %)

Some variations of the PiMS:

- CCUUCCCAGGUAU-AAACCAACC
- CCUCCCCAGGUAACAAACCAACC
- CCUUCCUAGGUAACAAACCAACC
- CCUCCCCAGGUAU-AAACCAACC
- CCUUCCCAGGUAACAAACCAUCC

### 4. Biophysical figures

**Figure 13, A**. Sequences analyzed. **B**, 1H NMR spectra and potential scheme of the i-Motif candidate examined in the 2019-nCoV. **C**, 1H NMR spectra of SARS-CoV version of CoVID-DNA.iM-1.

**A.**

| | | TAG | Type | Sequence | *in vitro* | Feature | Cons. | Other entries |
|---|---|---|---|---|---|---|---|---|
| PQS | 1 | CoVID-RNA.G4-1 | RNA | UGGCUGGCAAUGGCGGU | Y | N | 99.48 | Figure 2, A7; Figure 3, A6 &B2 |
| | 2 | CoVID-DNA.G4-1 | DNA | TGGCTGGCAATGGCGGT | Y | | | |
| | 3 | CoVID-RNA.G4-2 | RNA | UGGAGGAGGUGUUGCAGGA | Y | orf1ab - nsp3 | 99.78 | Figure 2, A4; Figure 3, B1 |
| PIMS | 4 | CoVID-RNA.iM-1 | RNA | AUCCUUCCUUACCCAGAUCCUA | N | orf1ab - nsp12 | 100 | Figure 2, A14; Figure 3, B4 |
| | 5 | CoVID-DNA.iM-1 | DNA | ATCCTTCCTTACCCAGATCCTA | Y | | | |
| | 6 | SARS-DNA.iM-1 | DNA | ATCCTGCCTTACCCAGATCCTA | N | | | Figure 3, B4 |

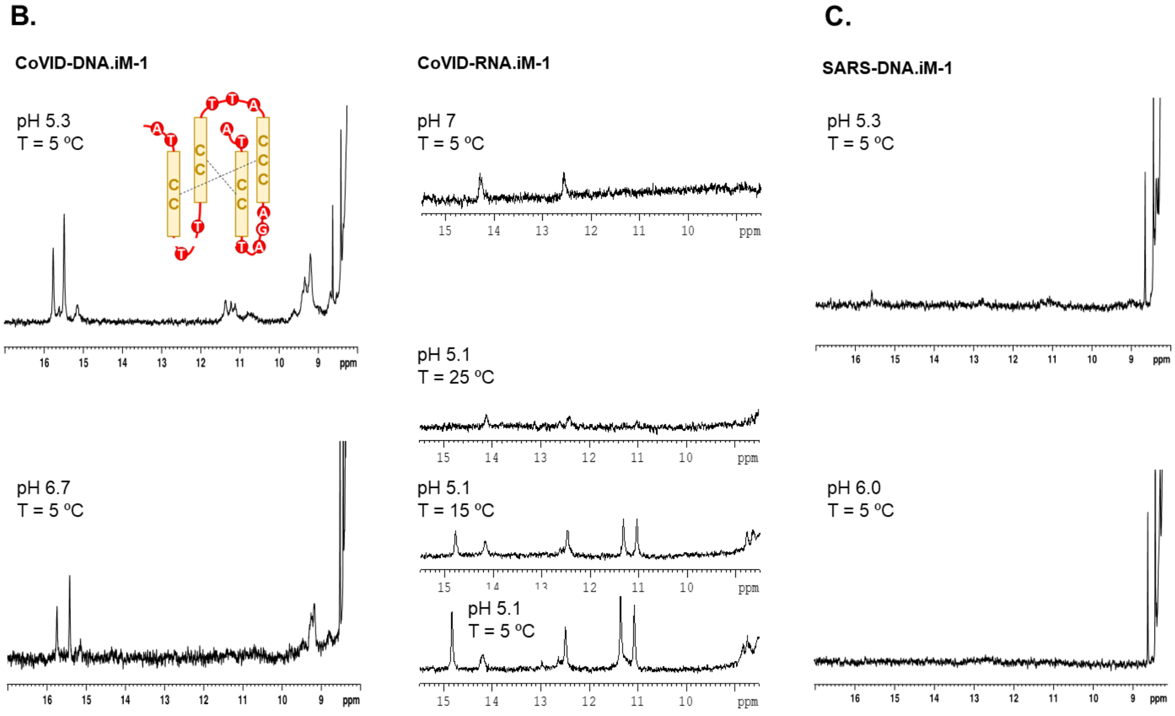

**Figure 14.** Regions of NOESY spectra of CoVID-DNA.iM-1 (left) and CoVID-RNA.iM-1 (right). Characteristic cross-peaks between imino (15-16 ppm) and amino (9.0-9.5 ppm) protons of ***C:C+*** base pairs are indicated on the left panels. The cross-peaks observed in the RNA are very weak and, most probably, correspond to AU and GC Watson-Crick base pairs. Spectra were recorded with 150 ms mixing time (25 mM sodium phosphate, pH 5.1, T = 5ºC).

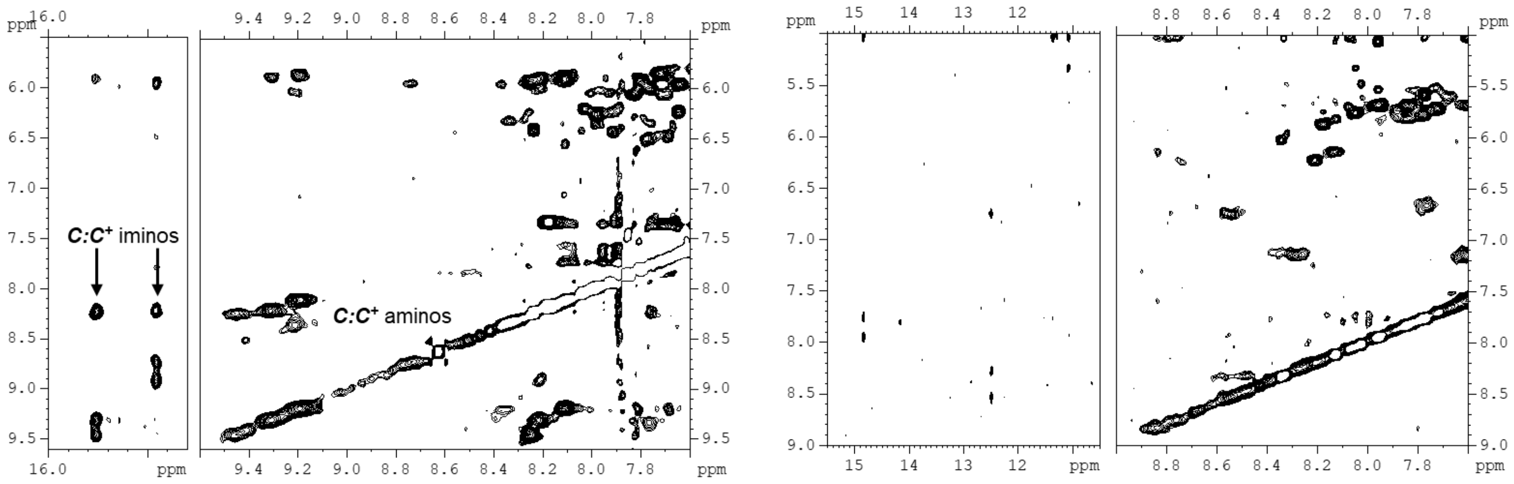

### 5. Data results

The data results for the entire analysis can be downloaded from the G4-iM Grinder's GitHub (https://github.com/EfresBR/G4iMGrinder). The package can be retrieved and installed also from here.

The data is divided into two groups.

#### G4-iM Grinder RAW data

*Virus.Results.RDS*, includes the raw data of the G4-iM Grinder analysis on all the virus realm as a list. The list groups virus species by their families. Each species list includes a PQS and PiMS sublist. These store the composition, location, known-quadruplex sequences presence and score (amongst others) of PQS/PiMS found in each virus. The information used in this analysis was Method 2; size restricted overlapping search method (PQSM2A data.frames), although Method 3 results are also included.

*3297.2019-nCoV.Results.RDS*, includes the raw data of the G4-iM Grinder analysis on all the 3297 different 2019-nCoV virus causers of clinical symptoms analysed in the work. These were used to calculate the conservation of each sequence found in the reference 2019-nCoV, and are given as a list. The list groups virus by strains. Each includes a PQS and PiMS sublist. These store the composition, location, known-quadruplex sequences presence and score (amongst others) of PQS/PiMS found in each virus. The information used in this analysis was Method 2; size restricted overlapping search method (PQSM2A data.frames), although Method 3 results are also included.

**G4-iM Grinder Analytical data**

*Analysis.RData*, is the analysis results on the raw G4-iM Grinder data. It includes 5 lists.

1) *a.Ref.2019nCoV* – Analysis with *G4iMGrinder* function of the GiG-package of the 2019-nCoV reference genome. It includes the Method 2 results of the reference genome with the conservation rates and common sequences found in other viruses. The biological landmarks affected by the candidates retrieved using the function *GiG.df.GenomicFeatures* are also stored here.
2) *Analysis.2019nCoV.3297genomes* – Analysis with *GiGList.Analysis* function of the GiG-package. 3297 genomes of the 2019-nCoV sequenced at different times and locations of the ongoing pandemic were examined. PQS and PiMS sublists are the analysis for PQS and PiMS respectively. *df.index* data frame stores the identification of each genome used.
3) *Analysis.Coronaviridae.fam* – Analysis with *GiGList.Analysis* function of the GiG-package of the *Coronaviridae* family. PQS and PiMS lists are the analysis for PQS and PiMS respectively. *df.index* data frame stores the identification of each genome used.
4) *Analysis.Virus.realm* - Analysis with *GiGList.Analysis* function of the GiG-package of the entire virus realm. PQS and PiMS lists are the analysis for PQS and PiMS respectively. *df.index* data frame stores the identification of each genome used. *Genome* data frame is the analysis with the function *GiG.Seq.Analysis*.
5) *Baltimore.C* – Baltimore Classification tables regarding each group characteristics and classification of each family into its group.

Additionally, a PDF is attached with the references of the genomes used in the 3297 genomes analyzed of the 2019-nCoV as given by the GISAID database. This archive is called *gisaid.3297._hcov-19.pdf*.
